# Evolution of brain-wide activity in the awake behaving mouse after acute fear by longitudinal manganese-enhanced MRI

**DOI:** 10.1101/2020.05.12.091389

**Authors:** Taylor W. Uselman, Daniel R. Barto, Russell E. Jacobs, Elaine L. Bearer

**Author notes:** Corresponding author; Elaine L. Bearer, MD-PhD, FAAAS, FCAP,; (ELB), (ELB), (ELB). These authors contributed equally (Taylor W. Uselman, Daniel R. Barto).

## Abstract

Life threatening fear after a single exposure evolves in a subset of vulnerable individuals to anxiety, which may persist for their lifetime. Yet neither the whole brain’s response to innate acute fear nor how brain activity evolves over time is known. Sustained neuronal activity may be a factor in the development of anxiety. We couple two experimental protocols to obtain a fear response leading to anxiety. Predator stress (PS) is a naturalistic approach that induces fear in rodents; and the serotonin transporter knockout (SERT-KO) mouse responds to PS with sustained defensive behavior. Behavior was monitored before, during and at short and long times after PS in WT and SERT-KO mice. Both genotypes responded to PS with defensive behavior, and SERT-KO retained defensive behavior for 23 days, while wild type (WT) mice return to baseline exploratory behavior by 9 days. Thus, differences in neural activity between WT and SERT-KO at 9 days after PS will identify neural correlates of persistent defensive behavior. We used longitudinal manganese-enhanced magnetic resonance imaging (MEMRI) to identify brain-wide neural activity between behavioral sessions. Mn^2+^ accumulation in active neurons occurs in awake behaving mice and is retrospectively imaged. To confirm expected effects of PS, behavior was monitored throughout. Following the same two cohorts of mice, WT and SERT-KO, longitudinally allowed unbiased quantitative comparisons of brain-wide activity by statistical parametric mapping (SPM). During natural behavior in WT, only low levels of activity-induced Mn^2+-^accumulation were detected, while much more accumulation appeared immediately after PS in both WT and SERT-KO, and evolved at 9 days to a new activity pattern at *p*<0.0001, *uncorr*., T=5.4. Patterns of accumulation differed between genotypes, with more regions of the brain and larger volumes within regions involved in SERT-KO than WT. A new computational segmentation analysis, using our *InVivo* Atlas based on a manganese-enhanced MR image of a living mouse, revealed dynamic changes in the volume of significantly enhanced voxels within each segment that differed between genotypes across 45 of 87 segmented regions. At Day 9 after PS, the striatum and ventral pallidum were active in both genotypes but more so in the anxious SERT-KO. SERT-KO also displayed sustained or increased volume of Mn^2+^ accumulation between Post-Fear and Day 9 in eight segments where activity was decreased or silenced in WT. Staining of the same mice fixed at the conclusion of imaging sessions for c-fos, a marker of neural activity, confirmed that MEMRI detected active neurons. Intensity measurements in 12 regions of interest (ROIs) supported the SPM results. Between group comparisons by SPM and of ROI measurements identified specific regions differing between time points and genotypes Thus we report brain-wide activity in response to a single exposure of acute fear, and, for the first time, its evolution to new activity patterns over time in individuals vulnerable to anxiety. Our results demonstrate the power of longitudinal MEMRI to discover how brain-wide activity evolves during recovery or persistence of fear responses leading to anxiety.

## Introduction

A single experience of life threatening fear induces lifelong anxiety in vulnerable people ^1^, yet the biological basis for this is unknown. Longitudinal MR imaging has the potential to reveal brain-wide activity changes accompanying the evolution of fear to an anxiety state. Since in human, capturing an acute life-threatening response is not feasible, we leveraged two conditions in mice to produce fear and image its evolution to anxiety: predator stress (PS) and knock-out of the serotonin transporter gene (SERT-KO) ^2^.

Predator stress can be induced by predator odor, a naturalistic stimulus, displayed as defensive behavior in rodents ^3-6^. For predator odor, the synthetic 2, 3, 5-trimethyl-3-thiazoline (TMT), a product of fox anal gland, can be measured and delivered in precise, reproducible amounts. TMT has been extensively used and validated for producing PS, in rodents, primates and even human ^7-12^ and for activation of neural activity in the olfactory system ^7,9^.

Progression from PS to anxiety is promoted by deletion of the serotonin transporter (SERT) gene ^13^. SERT-KO mice display prolonged defensive behavior after a single PS experience ^3^. SERT-KO mice are considered a validated “model” for post-traumatic stress disorder (PTSD) ^14^, which in humans is defined as a severe anxiety disorder arising after a life-threatening event ^15^. By contrasting brain-wide activity in WT with SERT-KO before, immediately and at long-term after PS, we expect to identify localized neural activities in response to acute innate fear and how those activities resolve, persist, or increase when evolving towards an anxiety state.

Mn^2+^-enhanced MR signal is a direct retrospective indicator of integrated neural activity occurring throughout the brain between systemic delivery of Mn^2+^ and subsequent MR scanning. Mn^2+^, a paramagnetic divalent cation, acts as a Ca^2+^analog, entering active neurons through voltage-gated Ca^2+^ channels, such as Cav1.2 ^16^, and is detectible as a hyper-intense signal in T_1_-weighted MRI ^17-19^. Thus MEMRI is unlike blood oxygen level dependent (BOLD) functional MRI (fMRI), which is a proxy for activity occurring in the scanner ^20^. BOLD fMRI is qualitatively different from MEMRI, relying on a complex dynamic between vascular and neuronal events. Both MEMRI and BOLD imaging are useful yet give different information. In small animal fMRI studies, presentation of the stimuli or behavioral events are typically performed within the scanner, and thus BOLD results may be limited by anesthesia, require adaptive training or restraint ^21,22^. fMRI requires fast imaging protocols to match the time dependence of BOLD events (seconds) and has the advantage of not requiring any exogenous agents. In contrast, Mn^2+^ enters active neurons over 24-48h after systemic delivery via tail vein, peritoneal cavity or subcutaneous tissue ^19,23-26^. In rat, signal may persist for 4 days and has dissipated by 14 days ^27^. Although radioactive Mn^2+^ may linger for months in the brain ^28,29^, this long-lived radioactive pool does not appear to be adequate to give a T_1_-weighted signal, or is not accessible for uptake in active neurons. Mn^2+^ delivered by intra-peritoneal injection (IP) accumulates more slowly in the brain than direct intravenous infusion ^30^. The hyper-intense signal at early time points likely represents both intra-neural Mn^2+^ and vascular/interstitial pools, since when mice are imaged at 24h and then exposed to new stimuli at 25-27h post-Mn^2+^ injection, new regions with Mn^2+-^enhancement appear ^31^. Thus a baseline image can be acquired, against which the consequences of stimulation can be detected.

MEMRI has been used to visualize functional activity maps at single times points in a number of species including mouse ^32-35^, rat ^36,37, Gobbo, 2012 #370,38-40^, avian ^41^ and non-human primate ^42,43^. In rat, MEMRI signal has been correlated with electrophysiology ^39^, and with intermediate early gene expression ^44^. MEMRI has been applied to understand patterns of neural activity in songbird vocalization ^45^; in neonatal mice for pain response ^46^and in rats for barrel whisker stimulation ^47,48^ and for predator odor response ^49^. In Aplysia, the sea slug, Mn^2+^ accumulation is proportional to excitatory neural activity measured by electrophysiology ^50,51^. Systemic Mn^2+^ does not abnormally activate neurons, as evidenced by lack of induction of intermediate early gene expression ^31^. Low dose systemic Mn^2+^ is non-toxic ^23,37,41^, and does not affect behavior even when perfused continuously for two weeks ^52^. Thus animals may be repeatedly injected IP ^42^, potentially producing rich MEMRI datasets to follow how activity evolves in awake behaving mice over time.

Here we follow the evolution of brain-wide neural activity from the freely behaving mouse, to immediately after a single PS exposure, then again at 9 days later to detect residual signal, and finally after a second IP Mn^2+^ injection to detect on-going and new activity (**Fig. 1, for more detail see Methods**). By performing a single fear exposure after capturing a “resting state” image, we compare brain-wide activity of freely exploring mice with their immediate fear responses, avoiding additional experience of another IP injection prior to the fear event. Behavioral measurements across all time points confirms the impact of TMT as a fear-stimulant, and the effect of SERT-KO on persistent defensive behavior. Between imaging sessions and during PS exposure and behavioral measures, mice are freely moving, during which time Mn^2+^ enters active neurons. We parse the resulting complex imaging data of 24 animals imaged at 5 time points by performing statistical parametric mapping followed by automated segmentation with our new high resolution *InVivo* Atlas of 87 regions, enabled by our skull-stripping and alignment programs ^53,54^. We then calculate the volume of activity in specific neuronal nuclei throughout the brain for each condition. Activity maps based on statistical significance are confirmed by region of interest analyses measuring intensity values, and further validated with c-fos histology. Results demonstrate a highly dynamic evolution of brain-wide activity during progression from fear to anxiety states dependent on the serotonergic system, and pinpoint specific brain regions involved.

**Figure 1:**
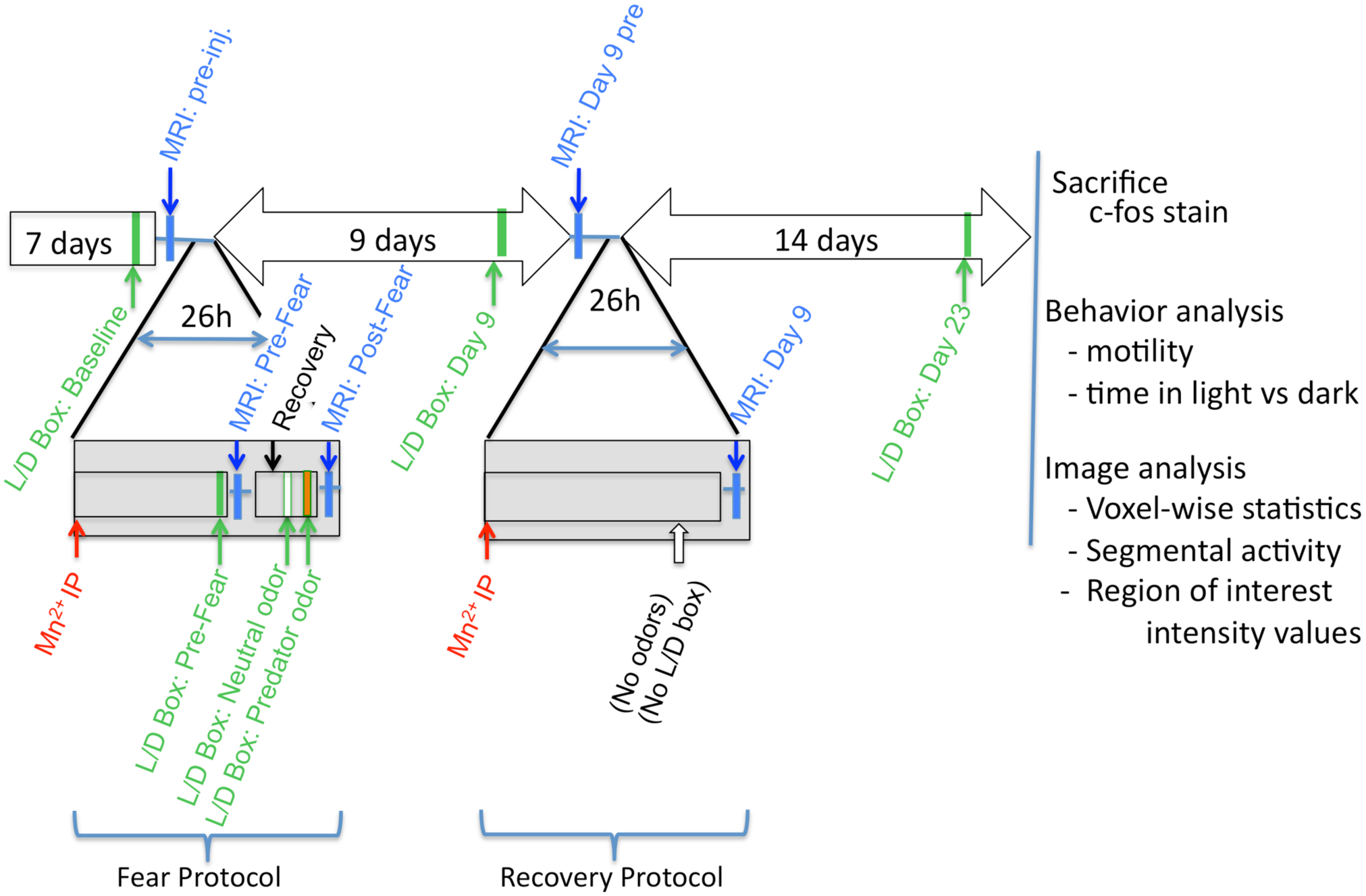
Schematic of experimental procedures. Open rectangles and block arrows indicate awake-behaving animals, horizontal single lines indicate anesthesia for imaging; Green arrows indicate behavioral recordings; Blue indicate MR imaging; Red indicate Mn^2+^ injection. Gray drop-down boxes are the two 26h experimental manipulations: Acute exposure to innate fear (“Fear Protocol”): Black bar, recovery from anesthesia; White bar, Neutral odor; Orange bar, Predator odor; Recovery or progression from acute fear (“Recovery Protocol”), with no second fear exposure; Open arrow, No odors, No behavioral recording. After arrival to the facility mice were acclimatized and handled daily for a minimum of 7 days. The experimental procedure began with a baseline behavioral recording in the light-dark box (L/D Box), and then capture of a pre-injection MR image. The “Fear Protocol” began with an intra-peritoneal Mn^2+^ injection after which mice were returned to the home cage for 22h for Mn^2+^ uptake to occur. Behavior was again recorded (L/D Box: Pre-Fear) followed by MR imaging (MRI: Pre-Fear). Afterwards mice were allowed ∼2h in the home cage to recover from anesthetic used in imaging (Recovery), and, when normally active, placed in the L/D Box again, first with saline (white bar L/D Box Neutral odor), followed by TMT, a predator odor (orange bar, L/D Box: Predator odor) for 30 min each, with behavior recorded in the last 10 min. Mice were then returned to the MR scanner (MRI: Post-Fear). After this image, mice returned to the home cage for 9 days after which behavior was recorded (L/D Box: Day 9), and a pre-injection MR image (MRI: Day 9 pre) captured to monitor any residual manganese-enhanced signal from the first Mn^2+^ injection. Mice then underwent another sequence, “Recovery Protocol,” which, except for the odors, was identical to the “Fear Protocol”--IP Mn^2+^, returned to the home cage for 24-26h, then imaged (MRI: Day 9), and returned to the home cage. At the conclusion of the procedure, mice are sacrificed and brains submitted for c-fos staining. Behavior and MR images are analyzed.

## Materials and Methods

### Ethics Statement

All protocols involving animals in this work were approved by Institutional Animal Care and Use Committees (IACUC) of California Institute of Technology and University of New Mexico. Animal numbers for all experiments were determined by a statistical power analysis based on pilot data. By performing experiments on the same group of animals over time, we obtain within group internal controls for each step in the procedure. Specific control experiments were performed on separate, smaller groups of animals to minimize animal numbers.

#### Animals

Mice were obtained from Jackson Laboratory (Jax.org, Bar Harbor, ME). Serotonin transporter (SERT-KO) knock-out mice (B6.129(Cg)-Slc6a4^tm1Kpl^/J, (Stock No: 08355), originally donated by Dr. Dennis Murphy ^55^, were back-crossed for 12 generations to be congenic with C57BL/6J (Stock No: 000664), the strain used as wild-type (WT). In addition to mice used for the full protocol, WT mice were also used in control experiments to determine Mn^2+^ kinetics, effects of IP Mn^2+^ and anesthesia on behavior, and to compare Mn^2+^ accumulation and c-fos expression in mice without PS to mice with PS.

Mice acclimated for one week after delivery from JAX and were handled daily by trained veterinarian staff technologists. All mice were housed at Caltech’s Beckman Institute Biological Imaging Facility in the MR scanner mouse house to allow for longitudinal imaging studies outside the animal care barrier. Housing, imaging, and behavioral testing were all done within the Small Animal Magnetic Resonance Imaging Center, which is environmentally controlled and sound-proofed. The mice were group-housed by littermates and set on a 12h light/dark cycle with standard bedding, and food and water provided ad libitum. Animal Care technologists regularly inspected mice for signs of injury or toxicity resulting from Mn^2+^ injections, anesthesia or imaging and found none. Mice were aged on average 81 days at the time of fear stimulus. We used all male mice for this study as our pilot studies in which we compared all time points between male and female mice revealed no significant differences, and using the same gender improved housing logistics.

#### Animal Procedure Overview

A schematic is shown in **Fig. 1**. The entire longitudinal procedure lasts 32 days--7 days to settle after shipping, 1 day for fear protocol, 9 days to recover, 1 day for recovery protocol, and another 14 days until final behavioral measure. Gray drop-down boxes represent the two experimental procedures, “Fear Protocol” and “Recovery Protocol,” separated by 9 days. Green arrows indicate behavioral recordings; blue arrows, MR imaging; and red arrows, the two IP Mn^2+^ injections. The odors, neutral and predator, were delivered once, in the “Fear Protocol”, as indicated. Open horizontal rectangles/block arrows indicate awake behaving animals, and single horizontal lines indicate brief anesthesia for MR imaging.

Mn^2+^ accumulation depends on the degree and duration of neural activity ^19,20,23,26,32,50,56^ and the extracellular concentration of Mn^2+^ in the brain ^19,24,57,58^. We reported that a neuro-active hallucinogen, delivered 25h after Mn^2+^ IP, results in significant increases in Mn^2+^-dependent intensity in specific locations at 27h compared to 24h pre-drug images of the same animals, or to non-treated animals at 27h (*p*<0.0001) ^31^. Thus a strong stimulus that increases neuronal activity delivered during the height of Mn^2+^ brain intensity results in enhanced Mn^2+^ signal detectable after time intervals as short as two hours. This timing allows us to obtain a pre-stimulus image of basal neural activity prior to stimulation, as recommended ^27^. If the biology allowed us to boost the Mn^2+^ concentration and increase time for uptake, we might see more signal. However, higher Mn^2+^ concentrations were lethal in SERT-KO mice. We also ruled out repeated injections to increase Mn^2+^ concentrations ^24^, as these are associated with elevated stress hormone, cortisol ^52,57^. Thus because of SERT-KO’s physical fragility and vulnerability to stress from handling, such further manipulations would further complicate interpretation of predator stress responses. Since we observed increased signal with PS delivered at 26-28h after IP Mn^2+^ in our pilot studies and in published work with drug stimulation ^31^, we adopted this experimental timing. This timing minimized manipulations and also had the benefit of allowing us to capture a Pre-Fear image shortly before PS exposure in the same animal rather than in a separate cohort, as has been done previously ^59^. By measuring behavior before and after MR imaging with anesthesia, we confirmed that this procedure did not affect behavior in wild type (**Fig. 2**). Behavior measurements in the SERT-KO revealed its vulnerability to all manipulations, as detailed further in **Figure 4** below. By acquiring video recordings of mouse behavior, we confirmed that mice had fully recovered from any post-anesthesia effect at the time of the “Fear Protocol”.

**Fig 2:**
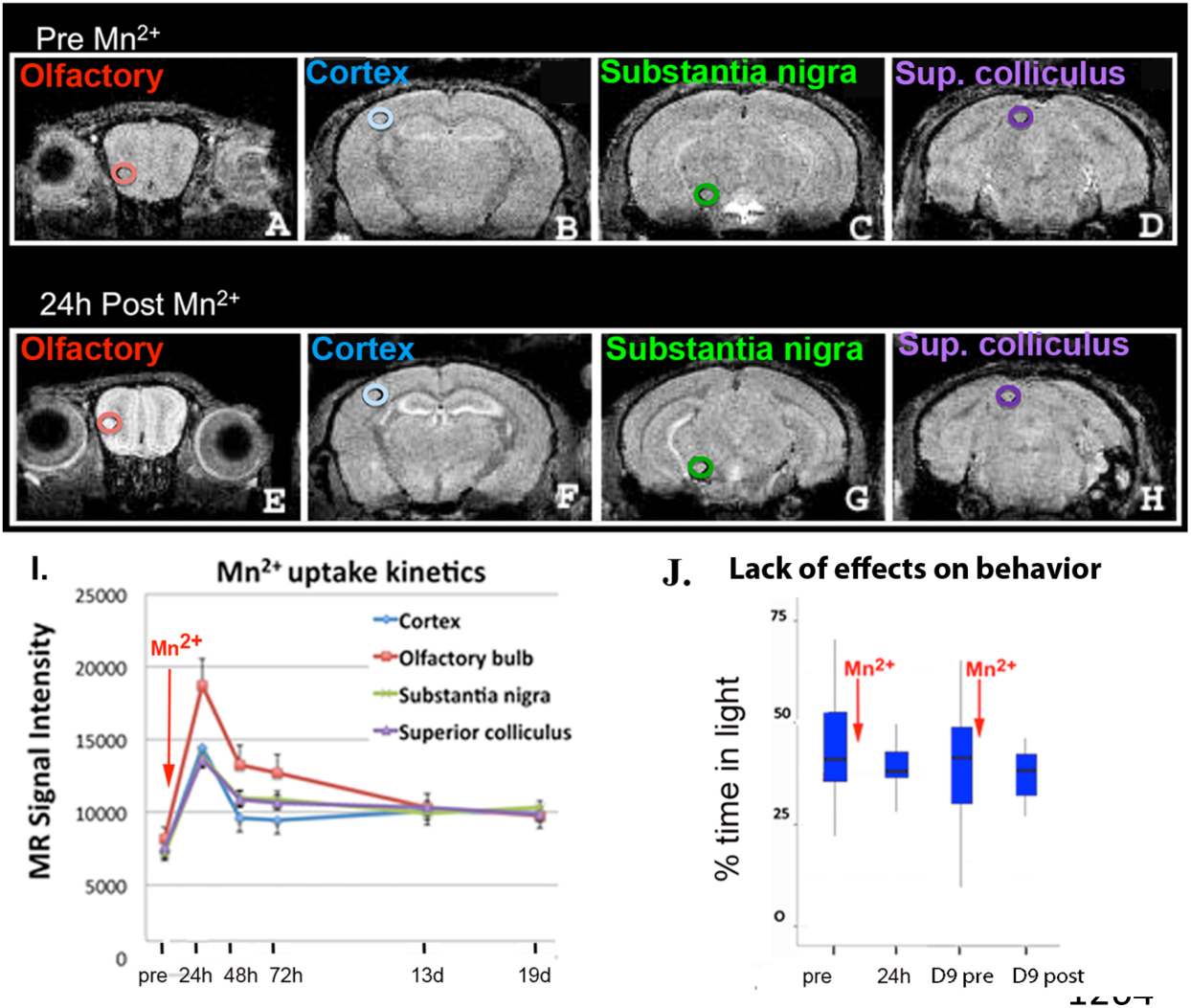
Mn^2+^ reaches a peak at 24 h after IP injection and the protocol without PS has no effect on WT behavior in the light-dark box. **A-H**) Representative examples of coronal slices before (**A-D**) and 24 h after IP Mn^2+^ injection (**E-H**). Locations where measurements were taken are indicated by circles, color-coded by region. MR pulse sequence: T2* FLASH, TR/TE 25/5ms, flip-angle 20 degrees, 100 micron isotropic voxels with total image time 34 min. **I**) Graph of intensity over time after Mn^2+^ injection, color-coded by region. Note that all four regions show a similar pattern. Also note that at 48h, 50% of intensity remains, and that even after 19 days intensity is still slightly above pre-injection. The rates of intensity increase likely represent a composite of absorption from the peritoneum into the vasculature, perfusion into the brain, entry into brain interstitium and accumulation in neurons; and likewise rates of decreased intensity would depend on excretion from the vasculature and clearance from the interstitium and neurons. **J**) Lack of effects on behavior of the protocol. Neither injection of Mn^2+^ nor the experience of anesthesia and imaging had an effect on average time spent in the light side of the box, even after a second IP injection and imaging 8 days after the first in WT mice without odor exposures (n=6). Quartile box-and-whisker plots with median indicated by horizontal bar inside the blue box, and 1st & 4th quartile distributions indicated by tails. There was no statistically significant difference in time spent in the light across time points (p=0.997), although a decrease in variance after Mn^2+^ injection and imaging was found, indicated by whiskers.

**Fig 3:**
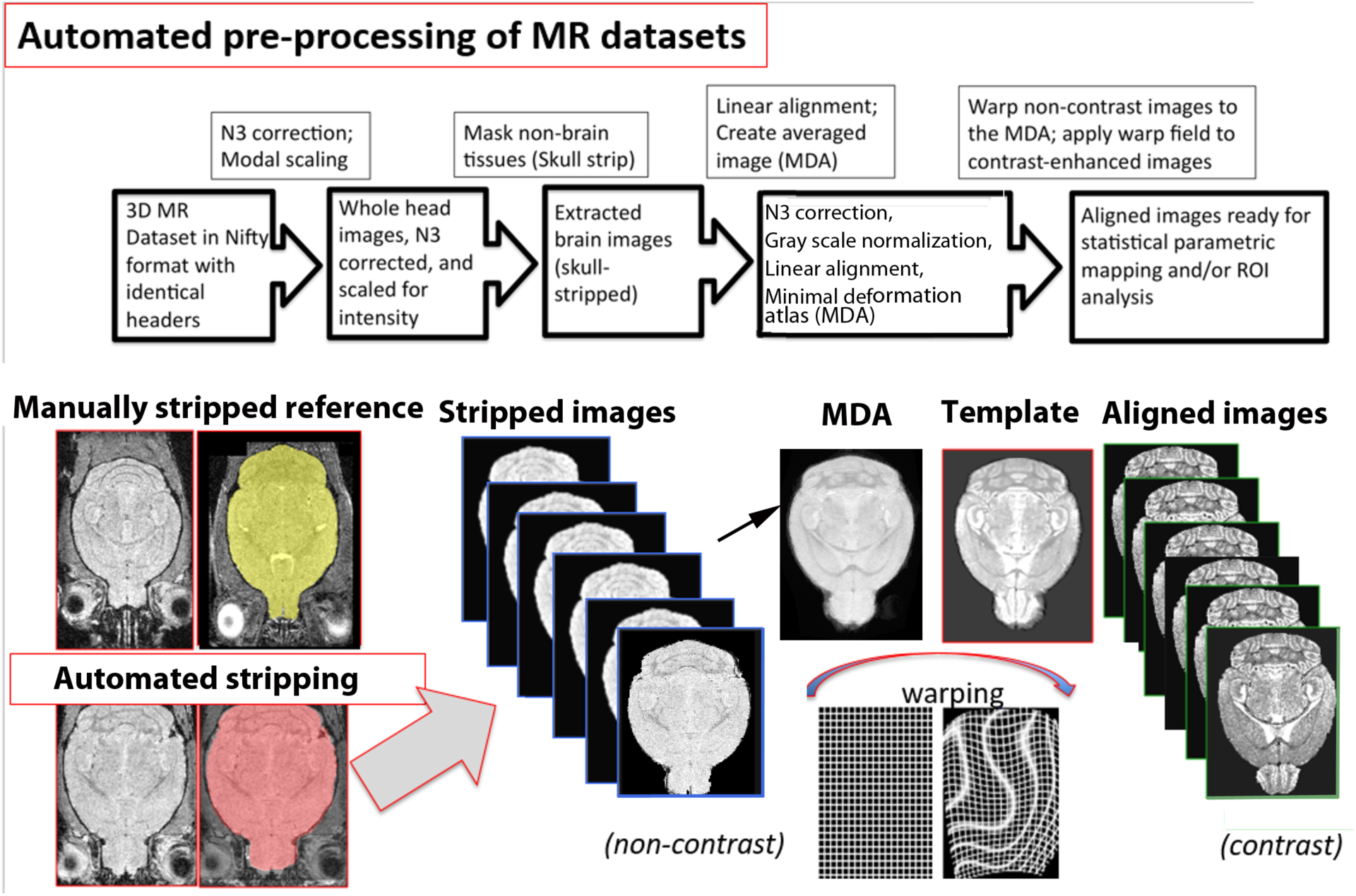
Diagram of automated processing steps: Skull stripping, registration and alignments. Flow chart of our automated pre-processing pipeline described in Methods (top panel), with examples of how images change with skull stripping and alignment steps (bottom panel). Raw images are brought into the same dimensions, resolution and image type in the header, then N3 correction is applied to equalize any inhomogeneities of the B field during scanning, followed by normalization of the intensity scale by modal scaling of the intensity histogram. Next the brain is extracted from whole head images with our automated skull-stripping software. Extracted brain images undergo a second N3 correction and modal scaling and then rigid body alignments. After linear alignment a minimal deformation atlas (MDA) is generated from the pre-injection, non-contrast-enhanced images. Each mouse’s non-contrast image is then warped (non-linear alignment) to the MDA and the warp field (control point grid) is applied to align the contrast-enhanced image for each mouse. The Mn^2+^-enhanced image (Template) used to create our *InVivo* Atlas is co-processed through all steps together with new data images. Align-warped images are ready for ROI analysis or to be smoothed for statistical parametric mapping. For examples of images at each step, see **Supporting Information Fig. S2**.

**Fig 4:**
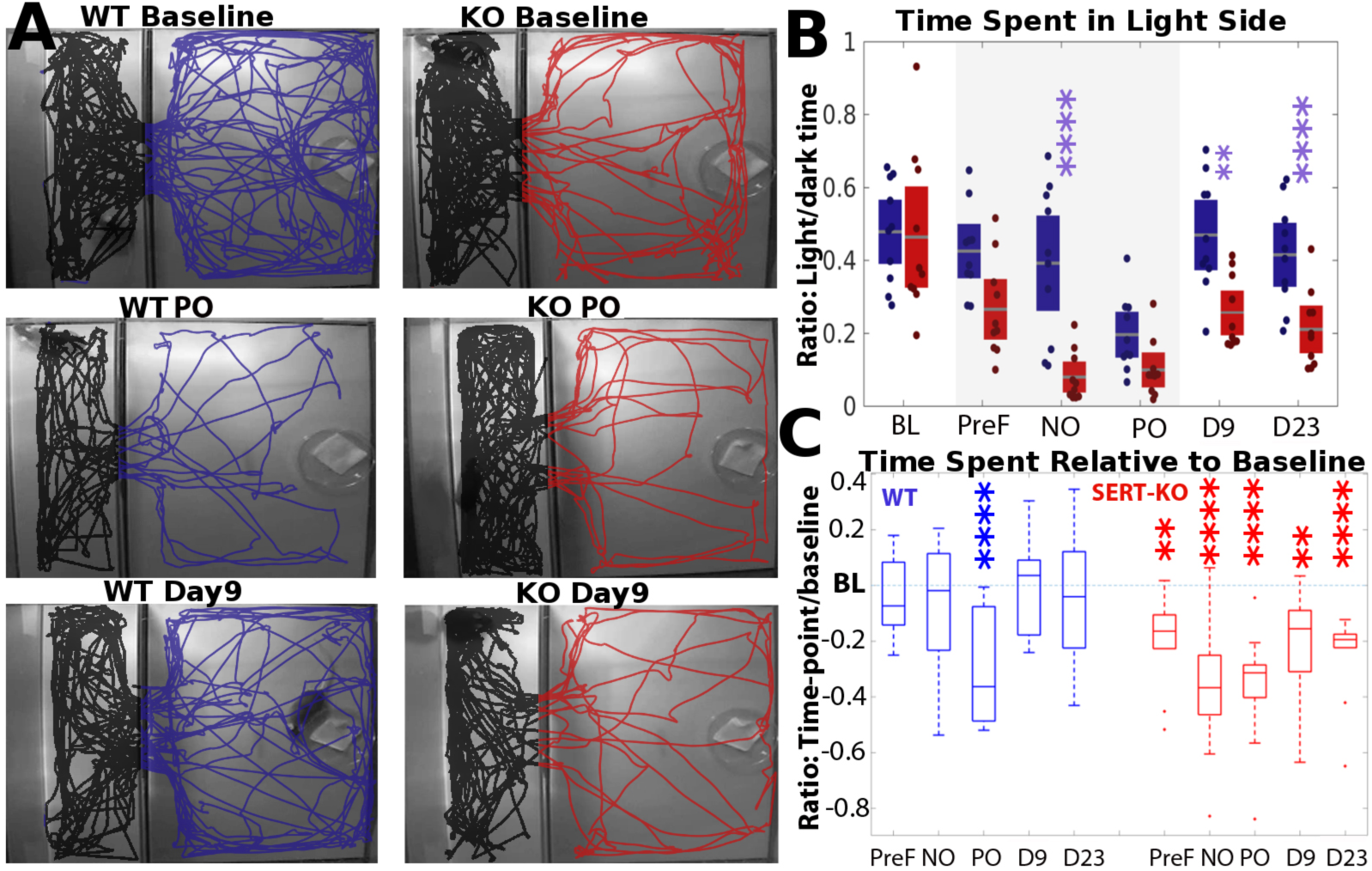
Time spent in the light before, during, and after a single predator stress exposure. **4A)** Examples of movement trajectories of representative WT mouse (blue) and SERT-KO mouse (red) over a 10 min session at Baseline (BL), Predator odor (PO) and Day 9 (D9). The light side of the box is to the right (colored tracings), and the dark side to the left (black tracings). Odors were introduced on a gauze pad in a petri dish visible on the lighted side. Note that the density of tracings, representing mouse motility, even during PO after imaging and anesthesia, appear similar to baseline, and only the location of trajectories is affected, whether in the light or dark side of the box. **4B)** Between group comparisons across time points shown as box and scatter plots, with WT (blue), SERT-KO (red). The gray line indicates the group average, with colored boxes +/- one SEM, and dots representing each individual. The shaded area indicates the three sessions acquired in the 26h “Fear Protocol”. Baseline (BL); Pre-Fear (PreF); neutral odor (NO); predator odor (PO); Day 9 (D9) and Day 23 (D23). ** *p*≤0.005; ****=*p*≤0.0001, 2-way ANOVA with Bonferroni correction. Note that WT mice displayed baseline exploratory behavior 24h, after Mn^2+^ (PreF) and before MR scanning, and also during neutral odor (NO), after the Pre-Fear scan that included anesthesia. This result demonstrated full recovery from anesthesia and no effect of scanning on this behavior. In contrast, SERT-KO already showed a decrease in ratio of time spent in the light at 24h post Mn^2+^ prior to the Pre-Fear scan, with further decrease during neutral odor presentation (NO). Also note that at Day 9 (D9) and Day 23 (D23), when no anesthesia or imaging had occurred, SERT-KO retained this defensive behavior. Hence the SERT-KO response to NO was likely a combination of all manipulations on this vulnerable genotype. **4C)** Within group comparisons of behavior at each time point (as in **B**), determined as change from Baseline (horizontal dotted blue line). Horizontal bars within each box plot represent the group median, the boxes are second and third quartiles, and whiskers are the range (minimum and maximum, first and fourth quartiles). Five blue boxes represent WT behavior and five red boxes SERT-KO. Dots indicate each individual animal, and those extending above and below the bars and whiskers represent outliers, defined by MATLAB boxplot function as 1.5 times the interquartile range between the 1^st^ and 3^rd^ quartiles. **=*p*≤0.01; \*\*\**p*≤0.001; ****=*p*≤0.0001 by ANOVA with post-hoc Dunnett correction. Note that there was no effect of Mn^2+^ injection or imaging experience, including anesthesia, on WT behavior. Only TMT had a significant effect. This lack of effect of neutral odor (NO) demonstrates that WT mice were sufficiently recovered from the imaging procedure and anesthesia at the time of behavioral testing. For SERT-KO, behavior at all time points after baseline displayed statistically significant decreases in time spent in the light, including at 9 days (D9), which was performed 9 days after the most recent anesthesia and imaging session, demonstrating that all manipulations had affected SERT-KO and arguing against any specific effect of delayed recovery from anesthesia.

Before any manipulations, a baseline of mouse behavior was recorded in the light dark box (L/D Box). A pre-injection MR image was then collected before administration of Mn^2+^. This image served as the non-contrast image to which all other images were compared. Next, each animal received an IP injection of Mn^2+^ for a whole body concentration of 0.3 mmoles/kg from a stock of warm 25mM MnCl_2_-4H_2_O solution in sterile saline. The injection volume was calculated based on the weight of each mouse, on average 26g.

At 22-24h after injection, behavior was recorded (L/D Box: Pre-Fear), and then mice were subjected to MR imaging (MRI: Pre-Fear). This first post-Mn^2+^ image represents an integrated picture of the mouse’s brain activity while freely moving in its cage in the environmentally controlled mouse house. After imaging, each mouse was awoken from anesthesia and allowed to recover for 2h in its home cage, then exposed in the light-dark box, first to saline, a neutral odor (L/D Box Neutral Odor), and then to predator odor, (L/D Box:

Predator odor), 2,3,5-Trimethyl-3-thiazoline (TMT) (ConTech, Victoria, BC, Canada Cat #v 300000368 lot 11125). We considered the entire Fear Protocol “predator stress” (PS). Each odorant exposure was for 30m, with Noldus video recording during the last 10m. Exposures were performed in a clean air hood to reduce external noise, smells or other sensory stimuli. Immediately after TMT, mice were again anesthetized and a second longer MR imaging session was performed. After the second imaging session, mice were returned to their home cages in the environmentally controlled sound-proofed mouse house. After 9 days, behavior was recorded (L/D Box: Day 9) and then a pre-injection MR scan was again captured (MRI: Day 9 pre). This scan monitored for any residual signal remaining from the first Mn^2+^ injection. Then each mouse was re-injected IP with Mn^2+^, returned to the home cage in the mouse house for 24h, and imaged again at 26h post-IP (MRI: Day 9), correlating with the same interval between Mn^2+^ IP and MRI as the Post-Fear scan (“Recovery Protocol”). Mice were returned to their home cage for 14 days and then behavior was recorded again in L/D box. Note that in each case the experimental condition occurred in the awake behaving animal, in a carefully controlled environment and that Mn^2+^ accumulation was detected retrospectively by MRI. Also note that there is only one fear experience--at 9 days there is no additional exposure. Finally, by recording with the Noldus video system, any residual anesthesia effects on motility could be rapidly detected, and none were found.

At the conclusion of the data collection, mice are sacrificed and perfused for histology, and behavioral recordings and images analyzed as described below.

#### Behavior

To monitor TMT-provoked defensive behavior, mouse activity was tracked in the L/D box with Noldus Ethovision (Ethovision XT9). The L/D box was a custom two-component plastic apparatus constructed by UNM Carpentry Shop and shipped to Caltech, with the dark side being a black walled enclosure (38 × 13 × 20 cm) with a dark tinted ceiling, or lid, that allowed infrared light to pass and a larger white-walled enclosure (38 × 28 × 20 cm) that had a translucent plastic ceiling that allowed illumination of the chamber. A floor level aperture (13 × 8 cm) allowed the mouse to move between compartments ^60^. The lid of the box contained odors within the box. Mice were randomly selected for testing regardless of genotype. The light-dark box was placed in a fume hood to reduce extraneous stimuli. Prior to behavioral recording after the Pre-Fear image, mice were recovered from anesthesia and allowed to explore their home cage for a minimum of 2h. Each mouse was placed in the dark-side chamber and left undisturbed for 30 min with a petri dish containing a gauze strip on the light side to acclimate to the chamber. For neutral odor, the gauze strip was first soaked in saline, and the mouse allowed to freely explore light and dark compartments for 30 min with recording during last 10 min. For TMT, the saline-soaked gauze strip was replaced with a gauze strip treated with TMT (600 µl of a 20 µl TMT mixed with 20 µl Ethanol and 582 µl of distilled water (3% solution)). Mice were again allowed to explore the box for 30 min with recording during last 10 min.

Mouse behavior was tracked by Ethovision ^61^ with the videos from all trials manually inspected to ensure no tracking errors, and that no mice had motor impairments, abnormal gait or evidence of residual anesthetic effects. After each trial, the box was wiped down with 10% ethanol to remove any residual odor. Only one mouse was tested in a single day, further reducing the chance of odor contamination. The fume hood cleared the air between trials.

#### Behavioral statistics

To quantify time spent in light versus dark side of the box, videos were analyzed using the Noldus Ethovision software which generates a .csv file. For statistics, we performed two-way repeated measures ANOVAs in GraphPad Prism (GraphPad Software, La Jolla California USA, www.graphpad.com). Between group factor was genotype and within subject factor were the six time points when behavior was recorded (see green bars in **Fig. 1**). Multiple comparisons between genotypes at all-time points were performed with post hoc Bonferroni correction ^62^. To determine whether mice returned to normal baseline exploration, an additional post hoc comparison was performed with Dunnett multiple comparison ^63^, which compares each time point to its baseline obtained prior to imaging and Mn^2+-^injection within each genotype.

### Manganese-enhanced magnetic resonance imaging (MEMRI)

To image retrospectively Mn^2+^ accumulation occurring in the awake behaving mouse, mice were imaged in an 11.7 Tesla 89 mm vertical bore Bruker BioSpin Avance DRX500 scanner (Bruker BioSpin Inc, Billerica, MA) equipped with a Micro2.5 gradient system and a 35mm linear birdcage radio frequency (RF) coil was used to acquire all mouse images ^53,54,64-72^. For *in vivo* imaging, mice were anesthetized with 1-2% isoflurane (vol/vol). Temperature and respiration were continuously monitored to remain within normal ranges of 37°C and 100-120/m respiration rate ^72^. To minimize movement, the animal’s head was secured in a Teflon® stereotaxic unit within the RF coil which also ensured reproducible placement of the mouse head within the magnet during imaging. We employed a fast low angle shot (FLASH) imaging sequence that gives a T_1_/T_2_ signal ^73^, 8 averages, TR/TE_eff_=25 ms/5 ms; matrix size of 200×124×82 and FOV 20.0 mm×12.4 mm×8.2 mm, yielding 100^3^ µm^3^ isotropic voxels. Five MR images were collected for each animal (**Fig. 1**). Imaging time was 34 min except after PS, when four 34 min images were collected and subsequently averaged, since the exposure time during which Mn^2+^ redistribution would occur was short (10 min). This averaging did not affect overall intensity measurements nor the pattern, as determined by statistical parametric mapping of single compared to averaged Post-Fear images (**Supporting Information Fig. S2**, and **Fig. S3**).

#### Mn^2+^ kinetics and effect on behavior

*<u>Kinetics of Mn</u>*^*<u>2+</u>*^ *<u>uptake in brain after IP injection</u>*<u>: The dose of Mn^2+^ was determined from a series of pilot experiments following recommendations (^24^ in which we found that greater than 0.3mmoles/kg IP was lethal. We previously reported that stimulation by a hallucinogen at 25hr post IP Mn^2+^ resulted in robust signal when imaged at 27h ^31^. To measure the kinetics of Mn^2+^ at longer time points, we performed a series of pilot experiments</u> by injecting 0.3mmoles/kg Mn^2+^ IP and imaging WT mice (n=5) at 24h, 48h, 72h, 13d and 19d with no odor exposure (**Fig. 2**). We also measured intensity in muscle outside the brain in the same images and found a similar time course of intensity changes (not shown).

Measurements show the highest brain intensity between 20 and 28h in four brain regions (**Fig. 2**) with only a 50% fall off by 48h that is sustained at 72h, and diminishes subsequently, while never completely returning to baseline even at 13 days.

We reasoned that intensity at 24h included extra-neuronal Mn^2+^, since signal increases at 24-27h with neurostimulation, indicating the availability of Mn^2+^ for uptake into active neurons. Inevitably, Mn^2+^ arrives more slowly into the brain after IP injection than after a tail-vein infusion, since absorption of ions and fluid from the peritoneal cavity into the vasculature is slow ^74,75^. Furthermore, Mn^2+^-induced intensity in the ventricles after IP injection lags behind that of tail-vein infusion^30^. This data forms the basis of our experimental procedure, with a 9-day pre-injection image to capture the pattern of residual Mn^2+^, followed by a second injection to replicate the conditions of the fear protocol, boosting the Mn^2+^ interstitial concentration for entry into any neurons that newly activated during progression to anxiety. Thus we can compare activity immediately after fear with that at 9 days.

##### No effect on exploratory behavior

To monitor the effect of IP injection and imaging with its anesthesia on behavior, WT mice (n=6) were monitored in the L/D box before and at 24h after Mn^2+^ injection and imaging, both initially and then again 9 days later.

At this dosage (0.3mmole/kg), neither Mn^2+^ IP nor the imaging experience had a statistically significant effect on average time spent in the light side of the L/D box, even after a second IP injection 9 days after the first, by repeated measures ANOVA (p=0.997) (**Fig. 2J**). A decreased variance in behavior was noted at 24h after each procedure, with complete recovery at 9 days. We also found that IP injection of higher amounts of MnCl_2_ (0.4 mmoles/kg) was lethal for SERT-KO. Lack of Mn^2+^ effect on behavior is consistent with other reports in rats using subcutaneous infusion over two weeks ^52^. These results confirmed that the procedure did not significantly affect behavior of WT.

### Preprocessing of MR images: Brain image extraction and image alignments

To perform statistical comparison, images need to be gray-scaled and aligned ^54,76^. All MR scans (five images of 24 mice for 120 whole-head images) were preprocessed using our protocols for brain extraction ^53^ and MR image alignment ^54^ (**Fig. 3**). Minimum deformation atlases (MDA) of the pre-injection images were generated as we described ^53,54^. The template image is available in from Medina et al. in that online Supplemental Materials^54^. Briefly, for this study, all alignments began by converting the Bruker Paravision images to NIFTI (.nii) format, preserving the voxel size of 0.1 × 0.1 × 0.1 mm ^77-80^ Image borders were cropped to 200×124×82 using the Medical Image Processing, Analysis and Visualization program (MIPAV) developed by NIH ^81^ such that all images had the same dimensions. Headers were checked to ensure identical headers.

Our group has pioneered protocols that standardize brain extraction such that extraneous tissue, present in live mouse imaging, does not interfere with brain alignments (**Fig. 3**) ^53,54^. To extract the brain (skull stripping), a rigid body transformation (6 degrees of freedom) was performed to align each mouse’s whole head image. These images were corrected for B-field inhomogeneity using the N3 method ^82^, followed by modal intensity scaling using our MATLAB script ^53^. Next a reference mask was manually generated using Brain Suite ^83^ based on one of the images in the dataset (reference whole head image). This drawn mask was then applied to all head images using our automated brain extraction protocol ^53^ with visual inspection of each stripped brain image to ensure accuracy. Any remaining non-brain tissue was removed manually in FSLeyes ^84^. The skull-stripped images were once again corrected for B-field inhomogeneity and the gray scale normalized by setting the mode of the histogram for all images to our template image ^54^. Stripped brains were then aligned with the normalization function in SPM ^85,86^ (**Fig. 3**). To validate our alignments we used visual inspection and CheckReg in SPM. Please refer to **Supporting Information Figs S2** for skull-stripped images before (**Fig. S2A**) and after (**Fig. S2B**) alignments.

#### Statistical parametric mapping

For statistical analysis using SPM ^87^(**Fig. 3**), aligned images were first smoothed with a full-width half-maximum (FWHM) Gaussian kernel, three times greater than the voxel size (0.3mm), as we previously described Delora, 2016 #41;Medina, 2017 #42}. To detect significant intensity changes due to accumulation of Mn^2+^ within genotypes, we performed a series of pairwise within group t-tests. Comparing post-Mn^2+^ time points with its pre-injection images for both genotypes at all time points produced a T_(1,11)_=5.4, *p*<0.0001, uncorrected with cluster size of 128. By requiring significant voxels be within a cluster, we eliminate false discovery of isolated random voxels, although by clustering we may miss signal in some small discrete, but relevant, nuclei of <128 voxels. To compare between groups, we performed two-sample between group t-tests in SPM between genotypes at each time point. We did not attempt a flexible factorial analysis, as the complexity of these factorial models makes them unable to accurately partition variance ^88,89^. Most likely one image, the SERT-KO post fear image, would be the factor accounting for the most variance. To ensure that our four-image averaging in the Post-Fear dataset did not introduce error by increasing detection uniquely for that time point, we also performed pair-wise SPM t-tests of a single image from the four captured Post-Fear compared with pre-injection images, and of the 4-image average with pre-injection images (**Supporting Information Fig. S3**). Very few statistically significant differences were found.

Videos of the statistical maps were rendered in Amira (Thermo-Fisher, https://www.thermofisher.com/us/en/home/industrial/electron-microscopy/electron-microscopy-instruments-workflow-solutions/3d-visualization-analysis-software/amira-life-sciences-biomedical.html) (**see Supporting Information Video S1**).

#### Segmentation Atlas

To parse the complex SPM results, we developed a digital segmentation atlas with 87 annotated regions based on a manganese-enhanced MR image of a living mouse brain with 80µm resolution ^90^. This segmentation atlas improves on others with fewer segments created from MR images of fixed brains with less contrast ^91^, and on those with more segments but lower resolution, which are available commercially (Ekam Solutions, Boston, MA). Our *InVivo* Atlas was hand-drawn, as we previously reported for a non-contrast enhanced image ^92^, based on visible gray-scale anatomy in our high contrast MR image with reference to histological atlases from Mouse Reference Atlas, Allen Institute ^93^ and Paxinos & Franklin ^94^.

Computational alignment with our 3D dataset was achieved by aligning the labeled atlas to our template image (aligned to our MDA) using FLIRT ^95,96^ in the FSL package ^97^ with a 12 parameter affine rotation, a mutual information cost function, and a nearest neighbor interpolation, which adjusts the image sizes between our dataset (100µm isotropic voxels) and the atlas (80µm isotropic voxels). Alignments were checked by visual inspection of overlays in FSLeyes ^84^.

#### Volumetric comparisons of significantly enhanced voxels

To quantify the total number of voxels in each segment, the atlas was applied to the template image and the number of voxels calculated in fslmaths and stats ^98^. Individual masks for each of the 87 sub-regions in the *InVivo* Atlas were generated through fslmaths ^97,98^ and applied to T-Maps identified by SPM. Voxels with statistically significant intensity increases (*p*<0.0001, *uncorr.*, T=5.45) that fell within each segmentation mask were quantified with fslstats ^97,98^. The number of significantly increased voxels in each segment was divided by the total number of voxels of that segment to obtain a ratio of “activated” voxels. These ratios were plotted as column graphs or heatmaps with a customized MATLAB script (MATLAB 2016b, The MathWorks, Inc., Natick, Massachusetts, United States) (**Supporting Information Fig. S4 and S5**).

#### Regions of interest (ROI) analysis

To measure the magnitude of intensity greater than pre-injection in regions showing significantly enhanced voxels in SPM analyses, region of interest (ROI) analyses were performed. ROI coordinates were selected based upon areas of statistically significant increase identified in paired t-tests contrasting WT Pre-Fear and WT Post-Fear images greater than WT pre-injection image. To compare between genotypes at Day 9, coordinates were selected based upon areas of strong signal in paired T-maps from each genotype at Day 9 greater than pre-injection projected onto the same gray scale template. These coordinates were propagated across all aligned, warped, unsmoothed images at Pre-Fear and Post-Fear of WT, and at Day 9 of both genotypes. The ROIs were generated by extracting a cube of 5×5×5 voxels at each selected coordinate, with a volume of 0.125mm^3^ centered on each location. The mean intensity and standard deviations of these sub-regions were calculated via fslstats ^95^. Two-way ANOVAs with FDR multiple comparison correction ^99^ were used to calculate statistical differences between conditions in each ROI using GraphPad Prism (GraphPad Software, La Jolla California USA, www.graphpad.com).

#### Immunohistochemistry

To detect neural activity by an alternative method, we stained histologic sections for the intermediate gene c-fos ^100^. At the completion of the final imaging session, 6 mice of each genotype were deeply anesthetized and subjected to intra-cardiac perfusion of warm heparinized saline followed by 4% buffered paraformaldehyde as previously described ^67-71^. After perfusion, the head was removed, tissue stripped down to the skull, and brains in the skull incubated overnight in fixative. Brains were popped from the skulls and re-fixed for 1-2 days before embedding in gelatin for sectioning. All 12 brains were embedded in the same block and serially sectioned in register at 35µm intervals. Sections were collected in a series of 24 cups such that each cup contained serial sections through the whole brain of all 12 animals at ∼0.8mm intervals. For non-PS-exposed controls, mice (12 each WT and SERT-KO) were injected with Mn^2+^, imaged 4 times in the scanner and embedded and sectioned at the same interval as the PS-exposed mice. Cups were stained with anti-c-fos immunohistochemistry by Neuroscience Associates (NSA) (https://www.neuroscienceassociates.com).

Microscopy of sections was performed on an Olympus microscope with a 5 Megapixel camera using CellSens software and prepared as figures in Photoshop.

## Results

### Defensive behavior occurred in both WT and SERT-KO mice during the “Fear Protocol”

The “Fear Protocol” elicited defensive behavior in WT and SERT-KO mice acutely, which persisted in the SERT-KO for at least 23 days. Videos at Baseline (BL, before IP Mn^2+^), Pre-Fear (PreF, 24h after IP Mn^2+^), during exposure to saline (Neutral odor, NO) and to TMT (Predator odor, PO), and finally at 9 and 23 days (D9 and D23) recorded mouse motility and time spent in the light or dark side of the L/D box (**Fig. 4**). Tracings of movements of two individual mice showed exploration in the light by each genotype at baseline (BL) (**Fig. 4A**, top row), decreased time in the light during predator odor (PO) without a decrease in overall motility (**Fig. 4A**, middle row). At Day 9, WT returned to the light side more often than SERT-KO (**Fig. 4A**, bottom row). Similar motility of both genotypes was observed at PreF, neutral odor, and Day 23, although relative time spent in the light differed. This comparable motility at all time points demonstrates successful recovery from low anesthesia levels used for the earlier imaging session prior to the behavioral assay.

Quantitative analysis of time spent in the light or dark for all 24 mice in the dataset found a significant main effect of time point (F _(5, 90)_ = 16.39, *p* <0.0001), a significant main effect of genotype (F_(1, 18)_=20.06, *p*≤0.0003) and a significant interaction of time point and genotype (F_(5, 90)_=3.51, *p*≤0.006). Two post-hoc comparisons identified significant differences at each time point between genotypes (**Fig. 4B**), and between each time point compared to baseline behavior within each genotype (**Fig. 4C**).

Baseline (BL) behavior, obtained prior to pre-injection imaging, was not statistically different between genotypes. During the second behavioral session (PreF), obtained one day after the first pre-injection MRI and Mn^2+^ injection, WT displayed time in the light similar to baseline, but SERT-KO spent reduced time in the light, although difference between genotypes was not statistical significant (**Fig. 4B**). During odor presentation (neutral odor, NO, and predator odor, PO), the vulnerable SERT-KO avoided the light side of the box even with neutral odor (NO) despite normal motility, while WT remained unaffected by NO (between genotypes, *p*<0.0001, Bonferroni corr.). With PO, WT and SERT-KO were not significantly different, with both groups displaying defensive behavior in the presence of TMT. WT recovered to baseline by Day 9 (D9) but SERT-KO did not (between genotypes, *p*≤0.005, Bonferroni corr.).

WT behavior was only statistically different from baseline during PO, while SERT-KO displayed increasing vulnerability beginning with the “Fear Protocol” (**Fig. 4C**). SERT-KO maintained statistically significant and persistent defensive behavior, even to D23 (within genotype difference from baseline at 23 days, *p* ≤0.0001). This defensive behavior of SERT-KO was likely a cumulative response to all manipulations. Thus the anxiety phenotype of this knock-out mouse represents a composite of frightening experiences and serotonergic deregulation. With PO, WT mice displayed defensive behavior not significantly different than SERT-KO. Without PS, there would have been no WT fear response and recovery to compare with SERT-KO.

### Neural activity paralleled behavioral measures

Differing patterns of brain-wide activity were identified by SPM corresponding to each condition for each genotype (*p*<0.0001 *uncorr*.T=5.4) (**Fig. 5**). In WT changes in signal in the olfactory bulb between Pre-Fear and Post-Fear confirmed an effect of TMT stimulation on Mn^2+^ accumulation during the 24-27h time frame with no signal prior to odor exposure in WT and a strong signal afterwards (**Fig. 5**, Wild type, Pre-Fear and Post-Fear, white arrows), correlating with normal behavior at PreF and defensive behavior during PO.

**Fig. 5:**
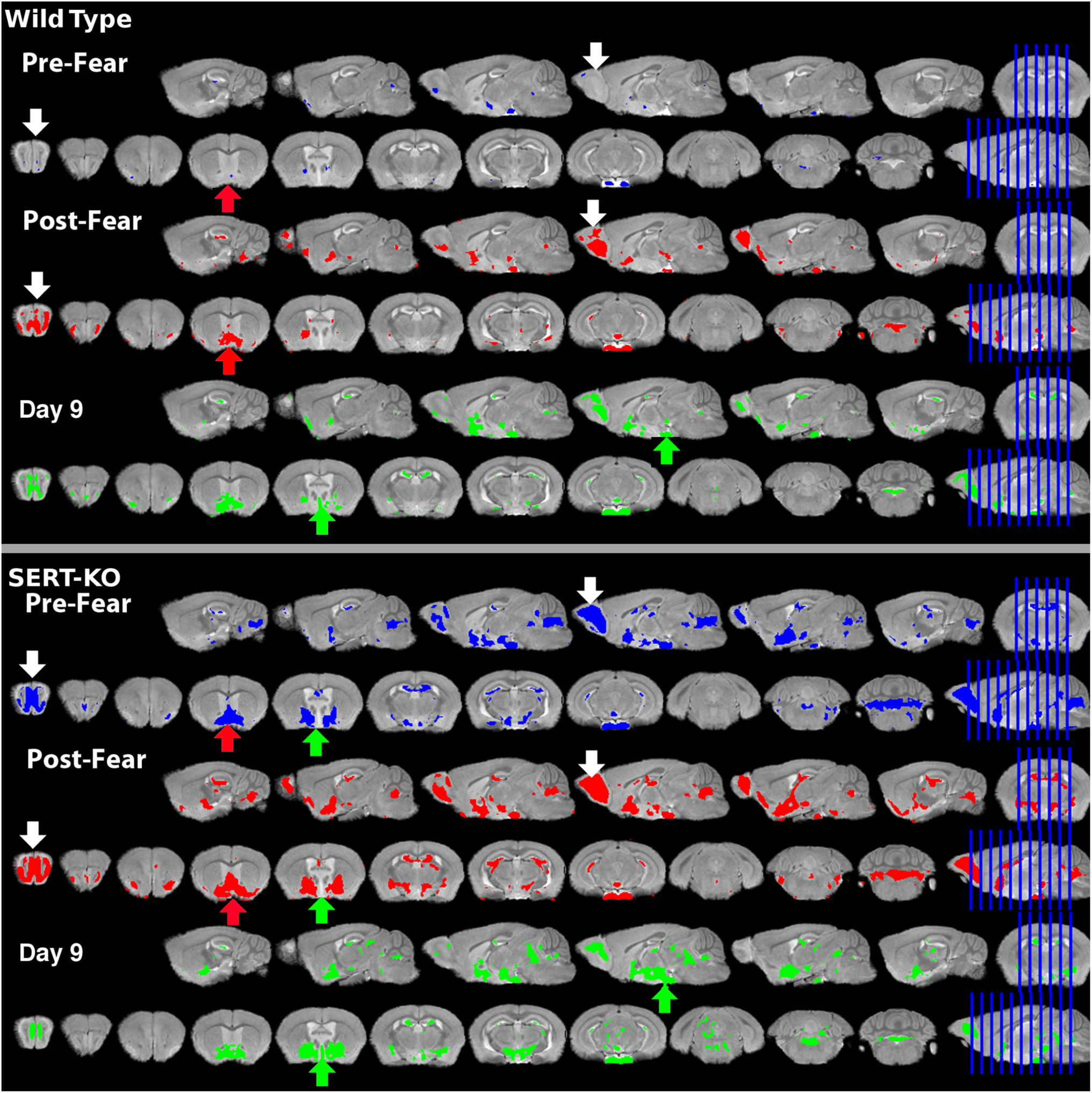
Statistical parametric maps (SPM) of Mn^2+^-induced intensity after each condition show statistically significant Mn^2+^ intensity increases in discrete locations across the brain. Shown is an overview of the imaging data, with a multi-slice panel of coronal and sagittal slices from six 3D SPMs projected onto the gray-scale MR template. Slices are evenly spaced across the 3D images in sagittal and coronal orientations, slice locations (rightmost images, blue lines). Top panel: Wild type, Pre-Fear (blue), Post-Fear (red), and Day 9 (green). Bottom panel: SERT-KO, similarly color-coded. Each condition was contrasted to the pre-injection images for that genotype using paired t-tests by SPM, *p*<0.0001 *uncorr*,, (T=5.4). White arrows, olfactory bulb; red arrows, basal forebrain; green arrows, hypothalamus. Note in WT that olfactory bulb is highlighted only after predator odor, as is basal forebrain; and that hypothalamus activates only at Day 9. In SERT-KO, these regions are already active Pre-Fear and their volume of activity appears to increase Post-Fear and to remain active at Day 9. Other areas of signal are also present throughout the panel. See **Supporting Information Video S1**.

Significantly, very little activity was detected in Pre-Fear images of WT. Pre-Fear imaging was performed after the mice had experienced two sessions in the light-dark box, an MR scan with anesthesia and restraint during scanning, and an IP Mn^2+^ injection (see **Fig. 1**). Hence, signal is not induced by these manipulations in WT, consistent with the lack of behavioral changes.

In WT Post-Fear, signals appeared in many areas in addition to olfactory system, including basal forebrain (**Fig. 5**, Wild Type, Post-Fear, red arrow). Finally at Day 9, the distribution of significant voxels changed further to include the hypothalamus (**Fig. 5**, Wild Type, Day 9, green arrows), although some activity had dissipated, that remaining appeared more caudal, consistent with the behavioral return to baseline.

In contrast, SERT-KO at Pre-Fear already showed signal in olfactory bulb which increased Post Fear (Fig. 5, SERT-KO, Pre-Fear and Post-Fear, white arrows). Fear-associated locations were also enhanced in the SERT-KO Pre-Fear, such as hypothalamus (**Fig. 5**, SERT-KO, Pre-Fear, green arrow) and basal forebrain (red arrow), consistent with its defensive behavior during PreF. Note that basal forebrain activity was ignited by PS in WT, but already active in SERT-KO at Pre-Fear (red arrows). This signal in SERT-KO Pre-Fear images likely represents its vulnerability to effects from all preceding manipulations excepting the odors. Without this pre-stimulus imaging, attribution of the effect of fear would be difficult. In SERT-KO Post-Fear, volumes of activity increased in locations already active in Pre-Fear, including olfactory bulb (white arrows) basal forebrain (red arrows) and hypothalamus (green arrows). At Day 9 in SERT-KO, with persistent defensive behavior, activity remained high in most regions with new activity appearing caudally.

Parsing differences in activity in these complex 3D whole brain T-maps with visual inspection is difficult. Hence, to evaluate this data quantitatively we developed a new computational method that segments 87 regions and measures their relative activity.

### *InVivo* Atlas-based segmentation revealed that volumes of activity-dependent Mn^2+^ intensity within segments is dynamic

To annotate sub-regions and digitally extract voxel-wise information from specific regions, we created an MRI-based *InVivo* Atlas that segments 87 neuronal regions at 80µm resolution throughout the brain of a living mouse (**Fig. 6).** The atlas was created based on our template image, a manganese-enhanced image of a living mouse brain (**Fig. 6A**) ^54^, and manually segmented based on recognizable gray scale contours with reference to histological atlases (**Fig. 6B**) ^93,94^. Although encompassing 87 distinct segments, this atlas is not as comprehensive as histological atlases of fixed mouse brain with some histologically defined regions not included. Our *InVivo* Atlas is particularly useful for this MR data as the template image was captured on the same MR scanner yet at higher resolution than our new dataset, which thus can be aligned without altering the new data at 100µm resolution (**Fig. 6C**).

**Fig 6:**
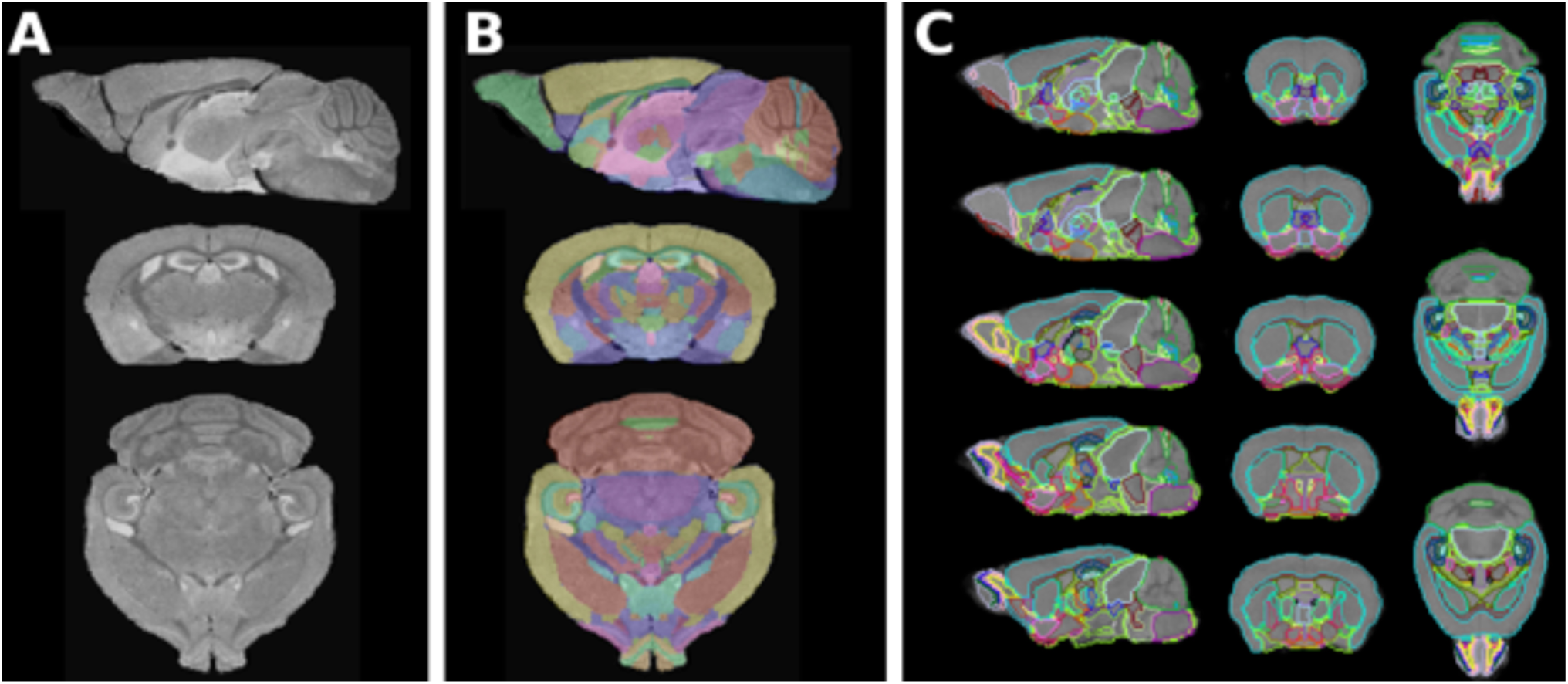
Segmentation by alignment of an *InVivo* MR atlas into this study-specific datasets. **A)** The template MEMR image of living mouse brain used to create the segmentation. **B)** Overlay of the *InVivo* atlas onto the template. Colored regions show an example of labeled segments. **C)** Overlay of the *InVivo* atlas onto the minimum deformation atlas from this dataset, shown in coronal, sagittal and axial slices with segments outlined in different colors.

Segmenting the T-maps allowed automated quantification of (1) numbers of voxels within each segment from the pre-injection anatomic image; and (2) numbers of voxels with significant intensity at specific threshold values from T-maps. With this information we calculate ratios of enhanced voxels versus total voxels for each segment at each time point for each genotype. Enhanced voxels were identified as significant at *p*<0.0001 *uncorr*. (T=5.4). We found 45 segments with ≥5% increase in the ratio of enhanced voxels compared to total voxels at any time across the datasets. SERT-KO displayed more significantly enhanced voxels than WT throughout the brain at all-time points.

How ratios of enhanced voxels change across 45 segments between pre-fear, post-fear and day 9 was visualized with column graphs and heat maps (**Figs. 7 and 8, and Supporting Information Fig. S4** and **S5**). Graphs, ordered by anatomy as columns and ranked by ratios in WT Pre-Fear images as heat maps, illustrate how regional activity evolves. Below we detail each condition/time-point in detail.

**Fig. 7:**
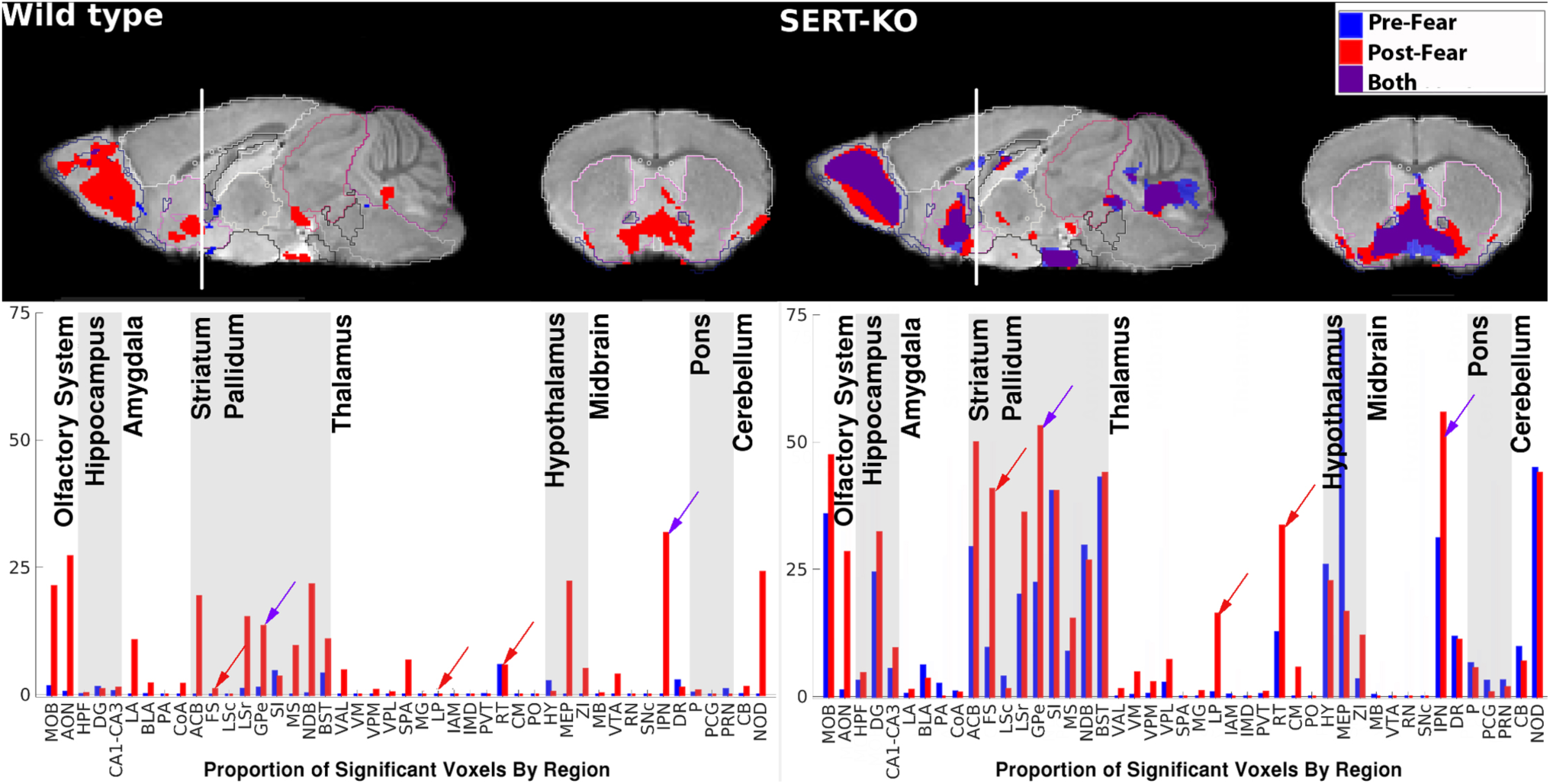
Identification and quantification of activity in 45 brain regions of Pre-and Post-Fear images for WT and SERT-KO. **Top Panel**: Sagittal and coronal overlay of Pre-Fear (blue), Post-Fear (red), and both (purple) (from bregma, sagittal: ML 0.18mm, coronal: AP 0.69mm) (*p*<0.0001 *uncorr*,, T=5.4). **Bottom panel**: Column graphs showing the ratio of enhanced/total voxels within segmented regions. Segments are listed on the x-axis from anterior to posterior (for abbreviations see **Table 1**). Purple arrows indicate examples of regions activated in both genotypes with predator odor. Red arrows indicate examples of regions with activity in SERT-KO and not WT after predator odor. Many other examples of increased activity in SERT-KO compared to WT are apparent (also see **Table 2**). For single Post-Fear images compared to averaged Post-Fear see **Supporting Information S3**. For complete array of column graphs including Day 9 pre, see **Supporting Information Fig. S4**. For heat map comparing ratios of enhanced/total voxels per segment, see **Supporting Information Fig. S5.**

**Fig. 8:**
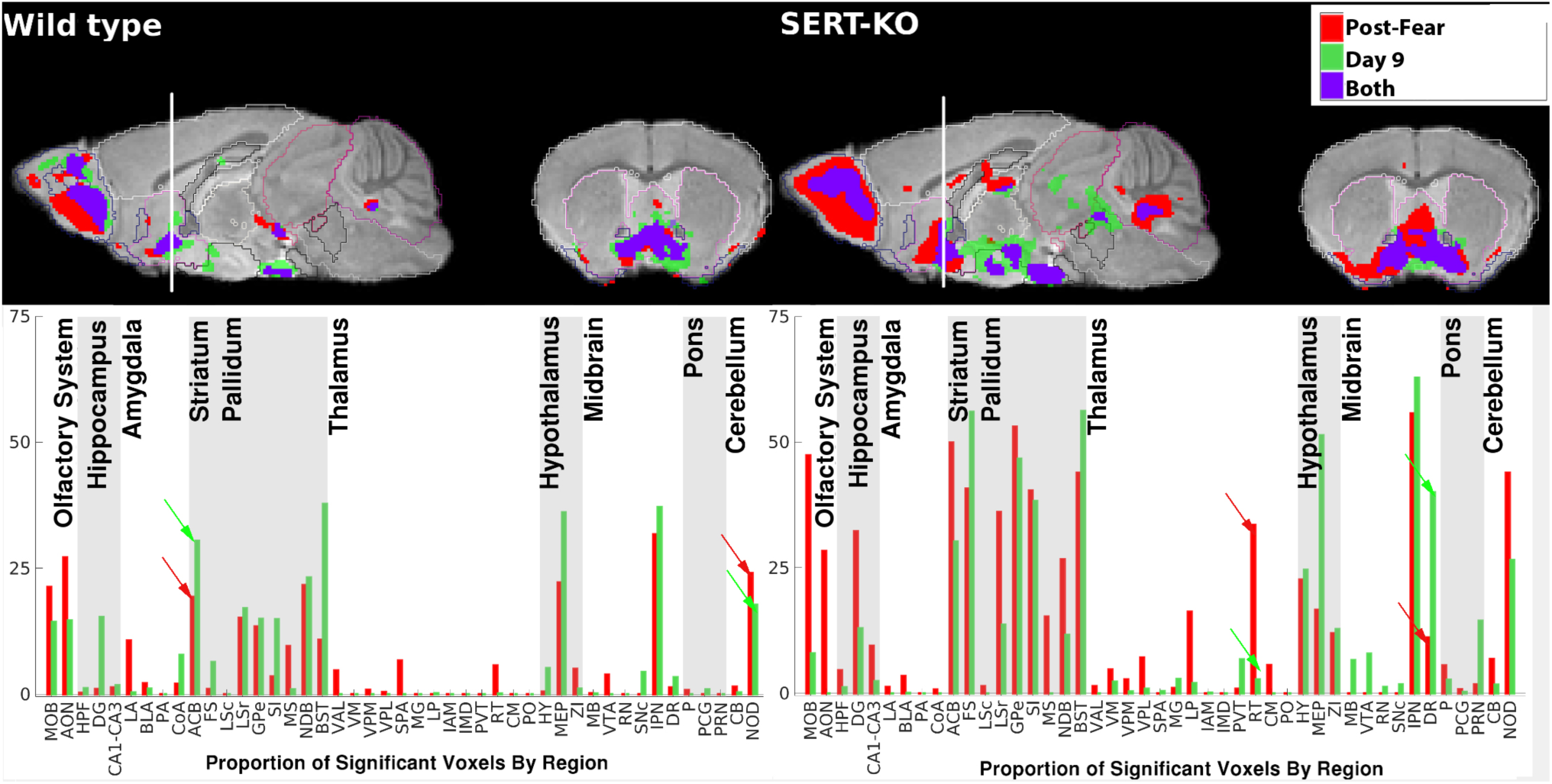
SERT-KO displayed prolonged neural activity at Day 9. **Top Panel:** Sagittal and coronal overlay of statistically significant voxels Post-Fear (red), and Day 9 (green), and both (purple). Note that there is no second predator odor after the “Fear protocol,” which occurred 9 days earlier. Note the additional Day 9 signal (green) in the SERT-KO sagittal slice (location of slices from bregma: sagittal slice: ML 0.18; coronal slice: AP 0.69mm) (*p*<0.0001 *uncorr*., T=5.4). **Bottom panel:** Column graphs showing activity within 45 segments that displayed >5% enhanced/total voxels in any image in the dataset (*p*<0.0001, T=5.4). Note high level of activity in the SERT-KO. Pairs of red and green arrows indicate examples of flips in amount of activity between regions from Post-Fear to Day 9. For example, in WT NOD has more enhanced voxels at Post-Fear than ACB, whereas at Day 9, ACB has more than NOD. In SERT-KO this flip appears between RT and DR, where RT decreases at Day 9 and DR increases. These changes in the balance of activity between segments can be found in other regions as well. For a heat map across time points, see **Supporting Information Fig. S5**; for complete array of column graphs, see **Supporting Information Fig. S4**. For between group comparisons at Day 9, see **Supporting Information Fig. S6. See Supporting Video S1.**

### A single exposure to acute fear (PS) resulted in increased Mn^2+^ accumulation in multiple regions of the brain compared to Pre-Fear

First we analyzed the differences between Pre- and Post-Fear in WT and in SERT-KO (**Fig. 7**, blue, Pre-Fear; red, Post-Fear, purple both). In Pre-Fear images, WT mice displayed few segments with activity ratios >5% whereas SERT-KO mice displayed activity ratios >25% in eleven segments, including the hypothalamus *(p*<0.0001 *uncorr.* T=5.4). Lack of signal in WT Pre-Fear (blue) could be because our statistical processing doesn’t detect individual variations in neural activity that would be expected in brains after naturally random exploratory behavior. In contrast in Post-Fear T-maps there is a group-wise effect with all mice responding more or less equivalently, and thus producing statistically significant changes. Increased neural activity in SERT-KO Pre-Fear may be due to its reported behavioral vulnerability to stress and possibly also to novelty ^3^, already manifest at PreF and worsening with NO in our behavioral testing. This significant increase in volume of enhanced voxels Post-Fear compared to Pre-Fear validates our experimental protocol, in which we first image a “resting state” pre-stimulus (Pre-Fear) at 24h post Mn^2+^ IP, and then stimulate with PS, imaging again immediately thereafter. Despite this short time frame for Mn^2+^ accumulation, significant additional signal is obtained.

In both WT and SERT-KO predator odor resulted in increased volume of enhancement of olfactory bulb (MOB) and anterior olfactory nucleus (AON) (**Fig. 7**, lower panels). Increased ratios in the olfactory system serve as an internal control that MEMRI has detected the expected TMT response ^44,49^. In WT, nineteen segments, including the olfactory system, increased signal in Post-Fear compared to Pre-Fear (*p*<0.0001, T=5.4) (**Fig. 7**, left panels, red columns). These increases were not due to averaging of four images captured Post-Fear, as single images gave virtually identical results (**Supporting Information Fig S3**).

In WT many segments displayed increases in the relative number of significant voxels between Pre- and Post-Fear in: amygdala, striatal structures (nucleus accumbens (ACB), bed nuclei of the stria terminalis (BST), lateral septum (LSr), medial septum (MS), and diagonal band (NDB)), thalamic nuclei involved in integrating activity between segments (ventral anterior lateral thalamic complex (VAL) and subparafascicular area (SPA)), hypothalamic and cerebellar structures (**Fig. 7**, left panels, see **Table 1 for Key to Abbreviations**). A small increase in enhancement was also detected in the ventral tegmental area (VTA). **Table 2** summarizes the changes between Pre-Fear and Post-Fear, ordered anatomically as in the column graphs and separated into two sections, one with those segments increased in WT Post-Fear (**Table 2A**) and the other with those increased in SERT-KO Post-Fear (**Table 2B**).

**Table 1.**
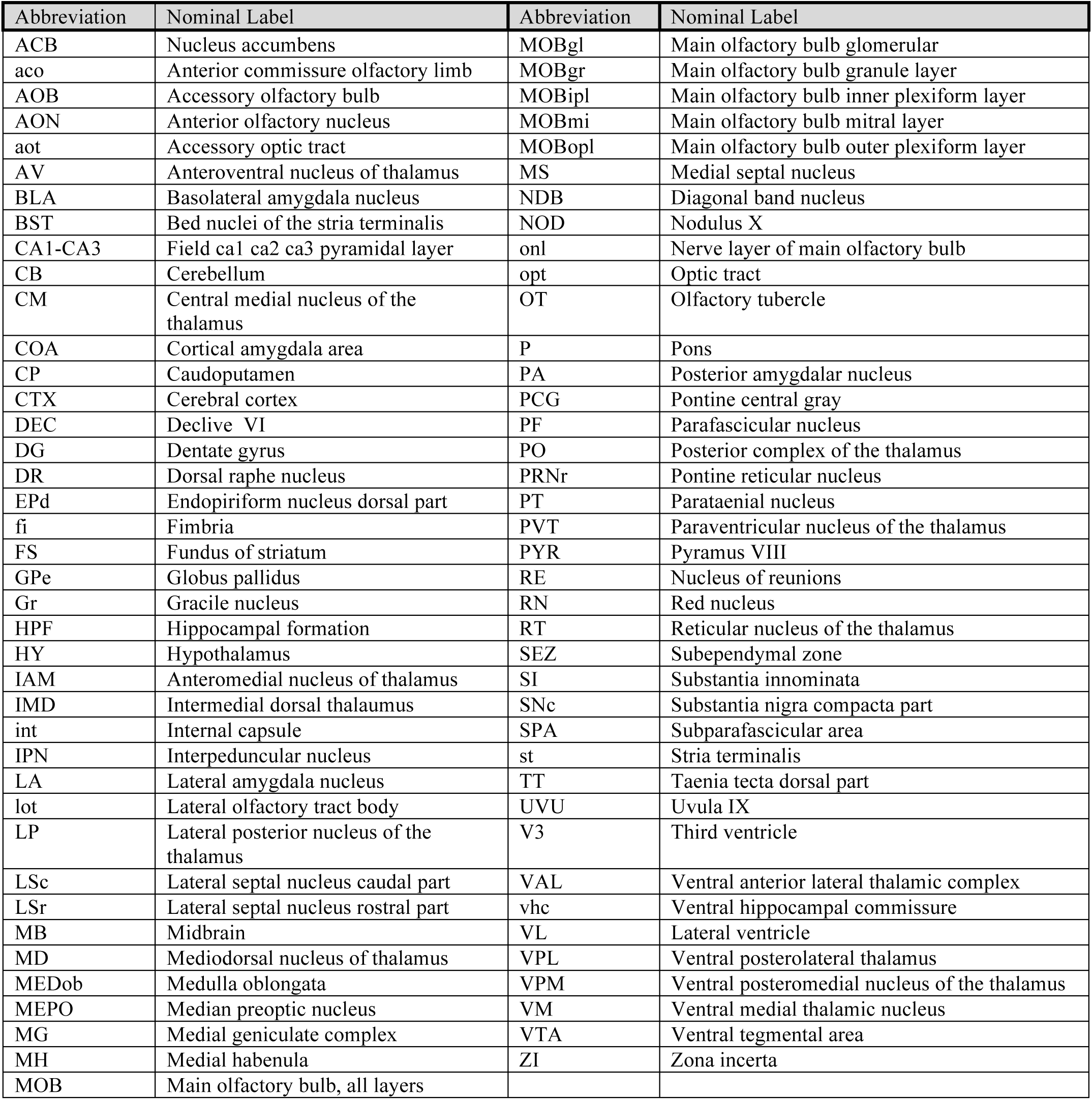
Key to abbreviations.

**Table 2:** Change between Pre-Fear and Post-Fear in WT and comparison with change SERT-KO mice.

| Abbrev. | Nominal label | WT | SERT |
| --- | --- | --- | --- |
| **A. Activity increased from Pre-Fear to Post-Fear in WT** |  |  |  |
| MOB & AON | Main olfactory bulb & Anterior olfactory nucleus | ++ | ++ |
| LA | Lateral Amygdala | ++ | + |
| BLA | Basolateral amygdala | + | - |
| COA | Cortical amygdala area | ++ | - |
| ACB | Nucleus accumbens | ++ | + |
| LSr | Lateral septal nucleus, rostral part | ++ | ++ |
| GPe | Globus pallidus | ++ | ++ |
| MS | Medial septum | ++ | ++ |
| NDB | Diagonal band nucleus | ++ | - |
| BST | Bed nucleus of the stria terminalis | ++ | = |
| SPA | Subparafascicular area | ++ | = |
| MEP | Medial preoptic area | ++ | -- |
| ZI | Zona incerta | ++ | ++ |
| VTA | Ventral tegmental area | ++ | N/A |
| IPN | Interpeduncular nucleus | ++ | ++ |
| CB | Cerebellum | + | - |
| NOD | Nodulus X | ++ | - |
| **B. Activity increased from Pre-Fear to Post-Fear in SERT** |  |  |  |
| DG | Dentate gyrus | = | ++ |
| CA1-CA3 | CA1-CA3 of hippocampus | = | + |
| FS | Fundus of striatum | = | ++ |
| LP | Lateral posterior nucleus of the thalamus | N/A | ++ |
| RT | Reticular nucleus of the thalamus | = | ++ |
| CM | Central medial nucleus of the thalamus | N/A | ++ |

In SERT-KO, Pre-Fear activity somewhat resembled WT Post-Fear (**Fig. 7**, right panel, blue columns). Purple arrows indicate examples of similarities, as if even before PO the SERT-KO brain was in a fear-like state corresponding to its decreased time spent in the light at PreF (**Fig. 4**). When the same segment in both genotypes increased activity in Post-Fear compared to Pre-Fear, the increase was greater in SERT-KO (**Fig. 7, both lower panels**). Even in Pre-Fear images, SERT-KO displayed high ratios of enhanced to total volumes in the main olfactory bulb (MOB) similar to those of WT Post-Fear.

In SERT-KO both Pre- and Post-Fear, high ratios were found in striatum and ventral pallidum (**Fig. 7**, bottom right panel). Many regions in SERT-KO that increased with fear were not affected in WT (**Fig. 7**, lower panels, examples indicated by red arrows). Many other examples of increased activity in SERT-KO compared to WT may be seen (**Table 2**). In contrast to WT, SERT-KO at both time points had no increased ratio in basolateral amygdala (BLA), but higher activity than WT in reticular nucleus of the thalamus (RT). SERT-KO also had strong activity in the hypothalamus Pre-Fear which was reduced Post-Fear (**Table 2**). Overall, SERT-KO had Pre-Fear activity patterns similar to a predator stressed WT, and this activity evolved differently than WT in a number of regions--six with increases and five with decreases (**Table 2**).

Although many of these segments are areas that Mn^2+^ seems to prefer ^25^, our statistical approaches reveal changes in them between conditions. Other than activity in cortical transition zones, cortical activity was not detected at *p*<0.0001, although MEMRI has been used to identify alterations of cortical activity in mutant mice ^101^ and cortical layer architecture ^102^. This failure to detect cortical activity was in part a result of our high statistical threshold of p<0.0001 (T=5.4), since at *p*<0.05 (T=1.8) bilateral activity is detectible by SPM in pre-limbic areas of medial prefrontal cortex and in retrosplenial cortex in Post-Fear SPMs of both genotypes (not shown). Our approach may under-estimate Post-Fear activity because of the relatively low dose of Mn^2+^, the timing of the PS and the short uptake period (2-3h). Despite this, we detected significantly increased activity in specific regions brain-wide in Post-Fear in both WT and SERT-KO.

### Brain activity evolved differently in SERT-KO from Post-Fear to Day 9 than in WT

To discover a brain-state corresponding to prolonged defensive behavior, we compared patterns of activity between WT and SERT-KO at Day 9 (**Fig. 8**, red, Post-Fear; green, Day 9; purple, both times; **Table 3**). We focused on Day 9 because at this time WT recovered baseline exploratory behavior while SERT-KO remained defensive ^3^, confirmed in our mice here (**Fig. 4**). Sustained defensive behavior in SERT-KO occurs in context of deregulation of the serotonergic system, and may represent a cumulative response of this vulnerable genotype to all experimental manipulations. Thus, SERT-KO provides a valuable opportunity to image an anxiety-related phenotype, which may not represent all types of anxiety produced by an acute fear event. WT, who returned to baseline behavior, revealed what normal recovery looks like, including complex transitions of continuing activity not manifested as defensive behavior in the L/D box. By comparing brain activity in SERT-KO at day 9 with WT, some anxiety-associated activity is revealed.

**Table 3:** Changes in activity from Post-Fear to Day 9, compared between genotypes.

| Abbrev. | Nominal label | WT | SERT |
| --- | --- | --- | --- |
| **A. Activity increased from Post-Fear to Day 9 in WT** |  |  |  |
| DG | Dentate gyrus | + | - |
| ACB | Nucleus accumbens | + | - |
| FS | Fundus of the striatum | + | + |
| SI | Substantia innominata | + | = |
| BST | Bed nucleus of the stria terminalis | + | + |
| MEP | Medial preoptic area | + | ++ |
| IPN | Interpeduncular nucleus | + | + |
| DR | Dorsal raphe | + | ++ |
| **B. Activity decreased from Post-Fear to Day 9 in WT** |  |  |  |
| MOB & AON | Main olfactory bulb & Anterior olfactory nucleus | - | - |
| LA | Lateral amygdala | - | = |
| MS | Medial septum | - | - |
| VAL | Ventral anterior lateral thalamic complex | - | = |
| SPA | Subparafascicular area | - | = |
| RT | Reticular nucleus of the thalamus | - | - |
| ZI | Zona incerta | - | = |
| VTA | Ventral tegmental area | - | + |
| NOD | Nodulus X | - | - |
| **C. Activity persisted from Post-Fear to Day 9 in WT** |  |  |  |
| LSr | Lateral septal nucleus caudal part | = | - |
| GPe | Globus pallidus | = | - |
| NDB | Diagonal band nucleus | = | - |
| **D. Activity increased from Post-Fear to Day 9 in SERT** |  |  |  |
| PVT | Paraventricular nucleus of the thalamus | N/A | + |
| MB | Midbrain | N/A | + |
| PRNr | Pontine reticular nucleus | N/A | ++ |

Each section (A-D) is ordered according to column graphs (Fig. 8)
+ >10% increase in activated ratio between Post-Fear and Day;
- >10% decrease in activated ratio between Post-Fear and Day;
= <10% change in ratio
N/A, no detected activity
Results are based on within genotype, paired t-tests in SPM ( $p < 0.0001$ , $T = 5.45$ ) as shown in Fig. 8.

To check for any residual signal from the first injection we captured a pre-injection image at Day 9 before injecting the second dose of Mn^2+^. We found minimal signal in this Day 9 pre image (See **Supporting Information Fig. S4** for column graph analyses). Thus the Day 9 post-Mn^2+^ signal represents new Mn^2+^-accumulation in neurons active then.

Brain-wide activity increased, decreased or persisted from Post-Fear to Day 9 in both genotypes (**Fig. 8, Table 3, and Supporting Information Fig. S5 and S6)**. We term these changes in activity over time “evolution of brain state,” which occurs spontaneously without any additional stimuli. More brain-wide activity at Day 9 was apparent in SERT-KO compared to WT. Activity decreased in both WT and SERT-KO in the olfactory bulb at Day 9 compared to Post-Fear, consistent with the absence of predator odor.

Our original hypothesis (**Supporting Information Fig. S1**) was that neural activity would resolve to Pre-Fear patterns in WT when behavior returned to baseline. Review of the WT imaging data comparing Post-Fear to Day 9 does not support this idea. In fact, activity increased in eight regions, decreased in nine and remained the same in three (**Fig. 8 and Table 3**). Increased activity at Day 9 occurred in four segments within the striatum and ventral pallidum (ACB, FS, SI, BST), as well as in hippocampus (DG), hypothalamus (MEP), and midbrain (IPN, DR) (**Fig. 8**, left panels, and **Table 3**). Thus, the picture is more complex, more nuanced, then we originally imagined.

Nine segments with decreased activity in WT at Day 9 compared to Post-Fear were: MOB, LA, MS, VAL, SPA, RT, ZI, VTA and NOD (**Table 3**). Activity persisted in three segments within the ventral pallidum: LSr, GPe, and NDB. This activity in WT at Day 9 was in mice whose exploratory behavior appeared grossly similar to baseline in the light/dark box, and thus may represent a memory of fearful experience that emerges on subsequent exposures and either provokes more dramatic responses or leads to resilience.

Segments with activity that decreased in both SERT-KO and WT at Day 9 are unlikely to contribute to defensive behavior, but rather may represent normal recovery processes. These include: MS, RT, and NOD (**Fig. 8** and **Table 3**). Segments with activity that persisted in WT and decreased in SERT-KO are also unlikely to contribute to defensive behavior, unless their function is inhibitory and silences abnormally active regions: LSr, GPe, NDB.

Segments at Day 9 in SERT-KO whose activity either persisted or increased in contrast to WT are most likely those that contribute to its prolonged defensive state. Three segments increased activity from Post-Fear to Day 9 in SERT-KO and were not active in WT at either time point: PVT, MB, and PRN. Segments that decreased in WT and persisted in SERT-KO included: LA, VAL, SPA, and ZI (**Table 3B**). The only segment that decreased activity in WT but increased in SERT-KO from Post-Fear to Day 9 was the VTA (**Table 3B**).

Analysis of segment-wise changes between Post-Fear images, when both groups are defensive, and Day 9 images, when WT recovers but not SERT-KO, presents one way to interpret this data. Alternatively, the balance of activity between segments across the brain may contribute to defensive behavior. For example, in WT NOD is more active than ACB at Post-Fear, whereas ACB is more active than NOD at Day 9 (**Fig. 8**, left panel, column graphs: Red arrows, Post-Fear; Green arrows, Day 9). Shifts in the balance of activity between segments is more dramatic in SERT-KO, with RT much more active than DR Post-Fear and the opposite at Day 9 (**Fig. 8**, right panel column graphs, red and green arrows). Finally, the degree of activity together with the balance of activity between segments may be the most significant in determining underlying brain activity correlating with defensive behavior. And there may be other regions, not yet detectible in these experiments, which also contribute to behavioral differences.

### Regions of interest (ROI) analyses supported statistical mapping, demonstrated magnitude of fear-induced neural activity changes, and quantified differences in the degree of activity between genotypes 9 days after fear

To quantify the magnitude of intensity changes, regions of interest (ROI) analyses were next performed. Six regions in WT were selected from Pre-Fear and Post-Fear image datasets based on statistically significant increases over pre-injection image by SPM (p<0.0001, *uncorr*., T= 5.4) **(Fig. 9A**). Significant differences of 3-8% between Pre-Fear and Post-Fear were found in all six regions: MOB, NDB, LSr, ACB, LA and SPA (*=p<0.05; **=p<0.01, *FDR*, by one-way ANOVA). Comparison of each of these regions by two way repeated measures ANOVA demonstrated a main effect of time F_(1,11)=_47.31, p<0.0001, with no significant main effect of segment. These results further document Mn^2+^ accumulation occurring with a stimulus at 24-27h after IP injection.

**Fig. 9:**
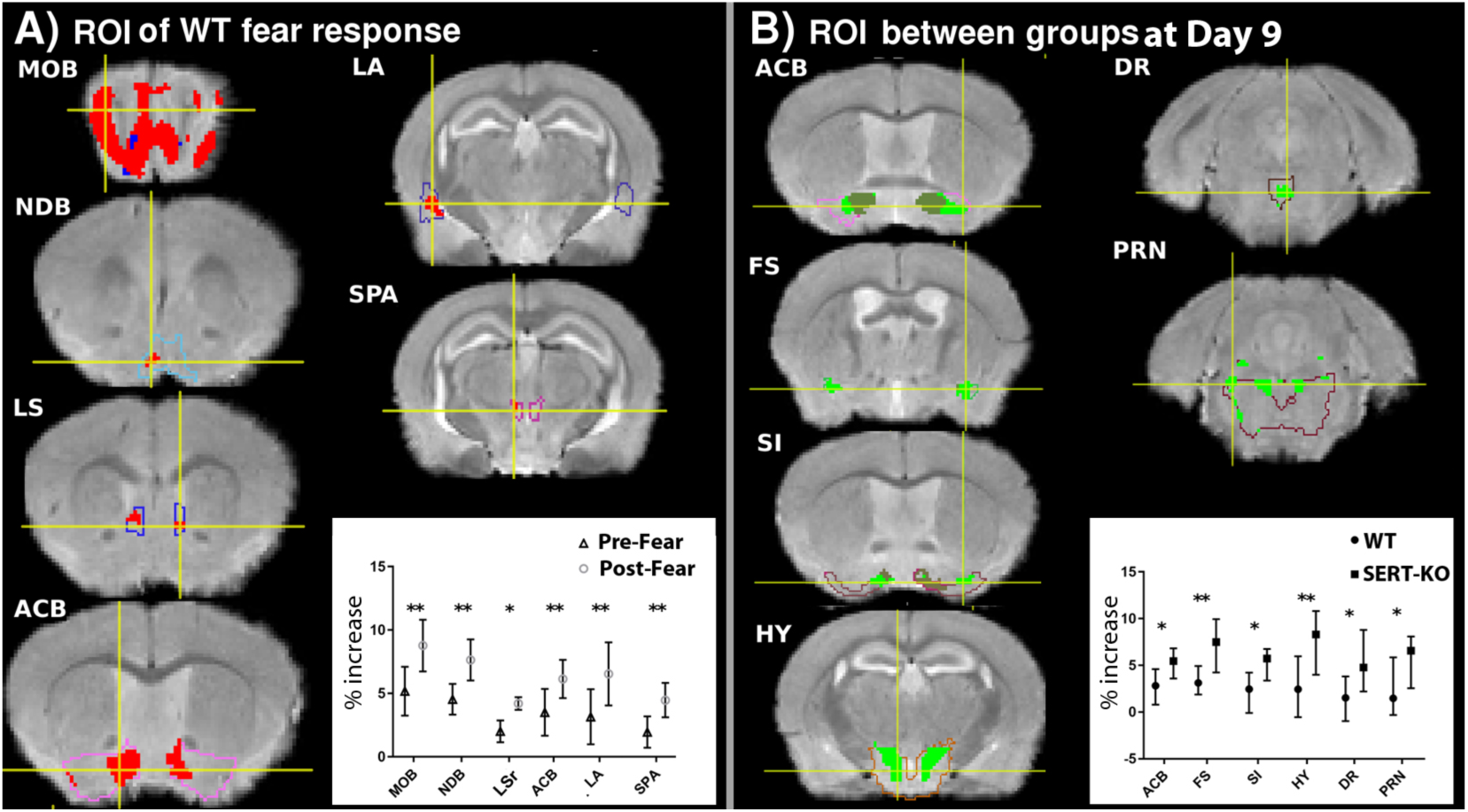
Magnitude of intensity between Pre-Fear and Post-Fear in WT validated SPM, and demonstrated that intensity differences between genotypes at Day 9 are significant. **A**) Comparison between Pre-Fear and Post-Fear intensity increase over pre-injection in six ROIs of WT (n=12), selected based on statistical maps, locations identified by alignment with the *InVivo* Atlas: Main olfactory bulb (MOB), diagonal band (NDB), lateral septum rostral region (LSr), nucleus accumbens (ACB), lateral amygdala (LA), and subparafascicular area (SPA). Coronal slices at the level of each segment are shown as labeled. Segments are outlined. Template (gray scale image) is overlaid with masked SPMs of that segment, with WT Pre-Fear (blue) and Post-Fear (red) (*p*<0.0001, *uncorr*. T=5.4). Minimal signal is detected at this statistical threshold in WT Pre-Fear. Yellow crossbars indicate the position of measurements, centered on 5 x 5 x 5 voxels. Graph inset: Percent of intensity increase of Post-Fear and Pre-Fear images over pre-injection image (vertical axis) for each segment at each time point: Pre-Fear (triangles) and Post-Fear (circles), with symbols at the mean and lines indicating +/- SEM. Differences in intensities between Pre- and Post-Fear are significant by repeated measures ANOVA with FDR correction. *=p<0.05, **=p<0.01, FDR. **B**) Comparison of intensity increase over pre-injection between genotypes at Day 9 in six ROIs selected based on statistically significant signal, locations identified by alignment with the *InVivo* Atlas. SPM images, with each segment masked, are overlaid onto the gray scale anatomic image, with WT in gold and SERT-KO in green. Green is unique SERT-KO activity and dark green is activity common to both genotypes where the two maps are superimposed. Segments are outlined. Slices are, from top to bottom: nucleus accumbens (ACB), fundus of striatum (FS), substantia innominata (SI), hypothalamus (HY), dorsal raphe (DR), and pontine reticular nucleus (PRN). Yellow crossbars indicate the position of measurements, centered on 5 x 5 x 5 voxels. Graph inset: Percent intensity increase over pre-injection of ROIs in Day 9 images for each genotype in each segment (vertical axis). WT (circles) and SERT-KO (squares) with symbols at the mean and lines indicating +/- SEM. Intensity values are significant using two-way ANOVA with FDR correction. *=p<0.05, **=p<0.01 FDR.

To compare differences between genotypes directly, we performed two between group analyses: (1) Comparison of intensity values in ROIs between Day 9 images of WT and SERT-KO (**Fig. 9B**); and (2) SPM unpaired t-tests between both genotypes at all time points (**Supporting Information Fig. S6**).

We selected six regions to measure in Day 9 images of both genotypes based on their significant intensity increase over pre-injection by SPM of SERT-KO at Day 9, and on their being previously implicated in fear and anxiety in other studies ^103^: ACB, SI, FS, HY, DR and PRN (**Fig. 9B, Table 1 and 3 for abbreviation keys**). Intensity values of Day 9 ranged from 7-11%, with SERT-KO values higher than WT in all ROIs. These differences in intensity values between genotypes were found statistically significant in all six regions (*=p<0.05; **=p<0.01 FDR by two-way ANOVA). Comparison of these measurements by two-way repeated measures ANOVA demonstrated a main effect of genotype F_(1,22)=_28.39, p<0.0001 with no significant effect of segment.

We also performed unpaired t-tests by SPM between genotypes at all three time-points for SERT-KO greater than WT, which gave signal at *p*<0.01 *uncorr.* (T=2.7). Results identified the FS, MG, PO, MB and PRN at Day 9 as having increased ratios of activated/total voxels in SERT-KO compared to WT (see **Table 1** for Abbreviation key, and **Supporting Information Fig. S6** for SPMs of un-paired t-tests and column graphs comparing segment-wise ratios of active to total voxels across all time-points between genotypes).

### Immediate Early Gene (IEG), c-fos, expression validates MEMRI

To further establish that SPMs of Mn^2+^-dependent intensity changes reflect neuronal activity, we performed c-fos immunohistochemistry as an alternative measure of brain-wide activity (**Fig. 10**). Subsets of brains from each genotype were fixed after conclusion of the protocol, embedded and sections stained for expression of the immediate early gene (IEG), c-fos, either with or without the Fear Protocol. Like Mn^2+^ uptake, c-fos expression is dependent on calcium channels that open upon neural excitation, and thus is an alternative indicator of active neurons ^104-106^. We previously reported that expression of another IEG, egr-1, is not induced by systemic Mn^2+^ or by anesthesia, and that its expression patterns matched MEMRI signal after psychopharmacologic treatment ^31^.

**Fig. 10:**
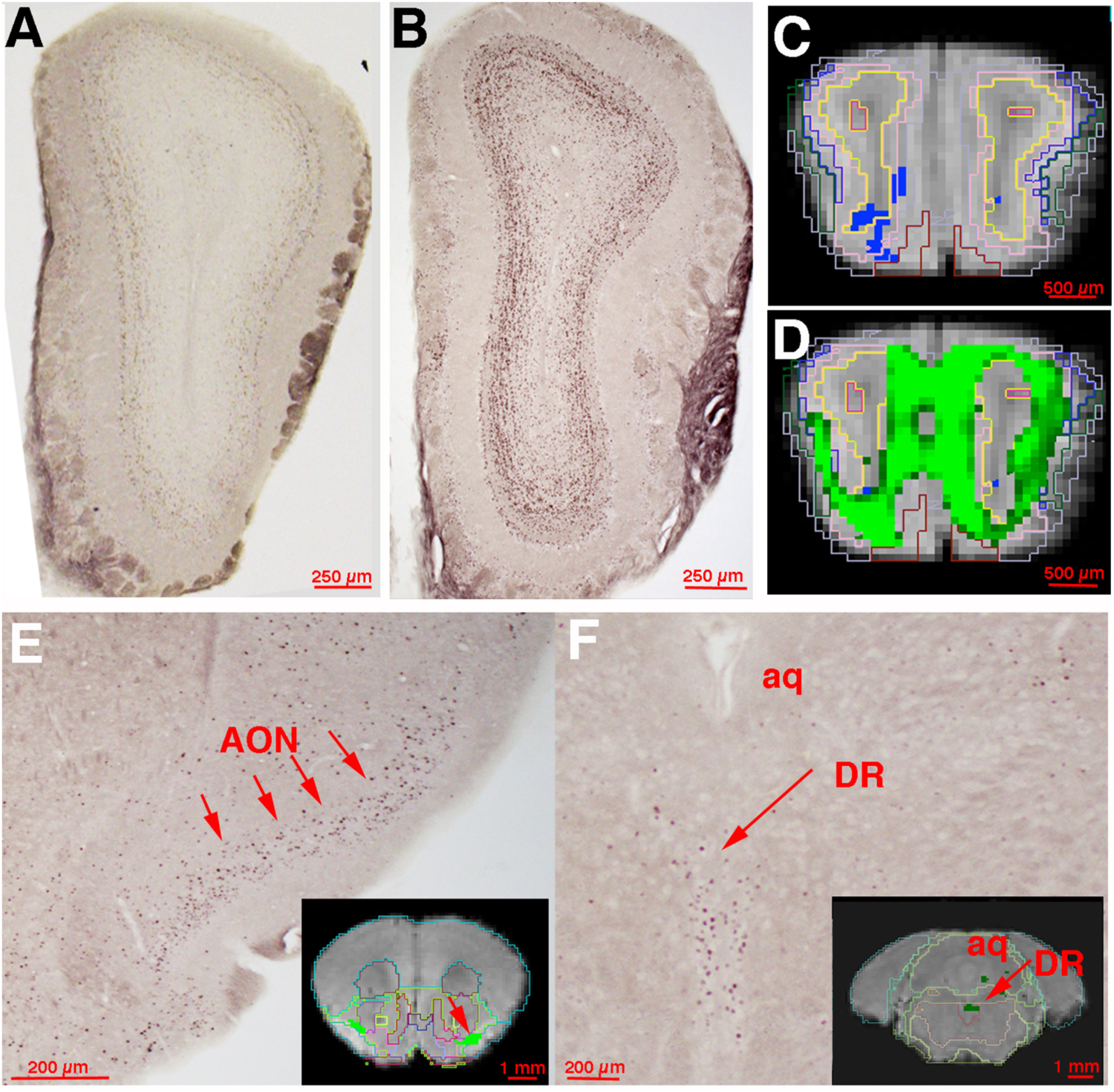
Immediate Early Gene (c-fos) expression validates the Mn^2+^ signal. **A-B**) Histological sections through the main olfactory bulb of WT mouse not exposed to TMT (**A**), and from a WT mouse that had experienced TMT (**B**). Note low level of c-fos stained nuclei in (**A**) and darker staining of the glomerular layer nuclei in (**B**) due to more nuclei stained for c-fos after TMT. **C)** SPM signal in the main olfactory bulb of Pre-Fear WT (blue) overlaid on gray scale template *(p*<0.0001 *uncorr.*, T=5.4), and **D)** SPM of main olfactory bulb at Day 9 in WT (green) overlaid on template (*p*<0.0001 *uncorr.*, T = 5.4). Note that neither c-fos nor MRI detects activity in anterior commissure olfactory limb or olfactory subependymal zone**. E)** Histological section through the anterior forebrain. Note signal (arrows) in anterior olfactory nucleus (AON) in both c-fos section and in MR image (inset, green SPM signal at D9 in WT) (*p*<0.0001 *uncorr.*, T=5.4). **F**) Histological section through the midbrain. The aquaduct (aq) provides orientation to the section location. The dorsal raphe nucleus (DR, arrow) is highlighted by both c-fos staining and by SPM of the MEMRI Day 9 (inset, green) (*p*<0.0001 *uncorr.*, T=5.4) surrounding areas are negative in both except for one small active area, very convincingly present in both c-fos and SPM images.

TMT-exposure induces c-fos expression in rats ^7-10^. The location of this expression correlates with MEMRI in the olfactory system, as we observe here (**Fig. 10A-D**). We examined WT and SERT-KO mice with and without TMT exposure and compared staining patterns in the olfactory system and dorsal raphe (DR). We observed strong c-fos nuclear staining in mice exposed to TMT in the olfactory system, including the main olfactory bulb (MOB) and anterior olfactory nucleus (AON), as well as the dorsal raphe (DR), concordant with SPM-identified Mn^+2^ enhanced intensity changes (**Fig. 10B-D**). Less staining was apparent in the non-TMT exposed mice that also had less MEMRI signal (**Fig. 10A** and **C**). Anatomic patterns in MOB of the TMT-exposed animals in both c-fos histology and MEMRI were strong in the granular layer, with a high density of nuclei, and less in the central subepidendymal zone. The AON and DR also displayed complimentary c-fos/MEMRI signal (**Fig. 10E** and **F**). Thus both positive and negative c-fos staining coincided with positive and negative MEMRI-based SPM signal anatomically, with or without TMT exposure.

MEMRI and immunohistochemistry are very different techniques, with different caveats and spatiotemporal dimensions. Preservation of c-fos activation is tricky and can be lost during fixation and processing. Our MEMRI images have 100µm resolution, while c-fos stained micrographs can be at the single cell level, or even as low as 500nm to 2µm. While we can set a threshold for signal significance in MEMRI, that is not yet possible with immunohistochemistry, where significance may be determined by numbers of stained nuclei within a certain area or volume rather than stain intensity. MEMRI is likely less sensitive than c-fos staining to low levels of neuronal activity, as is behavior. Hence the close similarity we show here between these two techniques, MEMRI and c-fos, is surprising and reassuring. A next step will be to acquire whole brain c-fos activation with such approaches as iDISCO and laser light sheet microscopy ^107,108^ and then to align resulting 3D IEG-stained images with MEMRI datasets ^109^, which will provide more precise information about the similarities and dissimilarities between these two techniques.

### Many enhanced regions crossed segment boundaries and some signal occurred in areas not annotated

T-maps did not respect segmental boundaries with highly enhanced segments contiguous at p<0.0001, T=5.4 (**Fig. 11**). This effect was particularly noticeable in the ventral pallidum of the SERT-KO Post-Fear and D9 images when statistically significant enhancement was most pronounced. These results suggest that parsing the brain into histologically distinct segments tells only part of the story--these segments may act together functionally in as yet undefined ways.

**Fig. 11:**
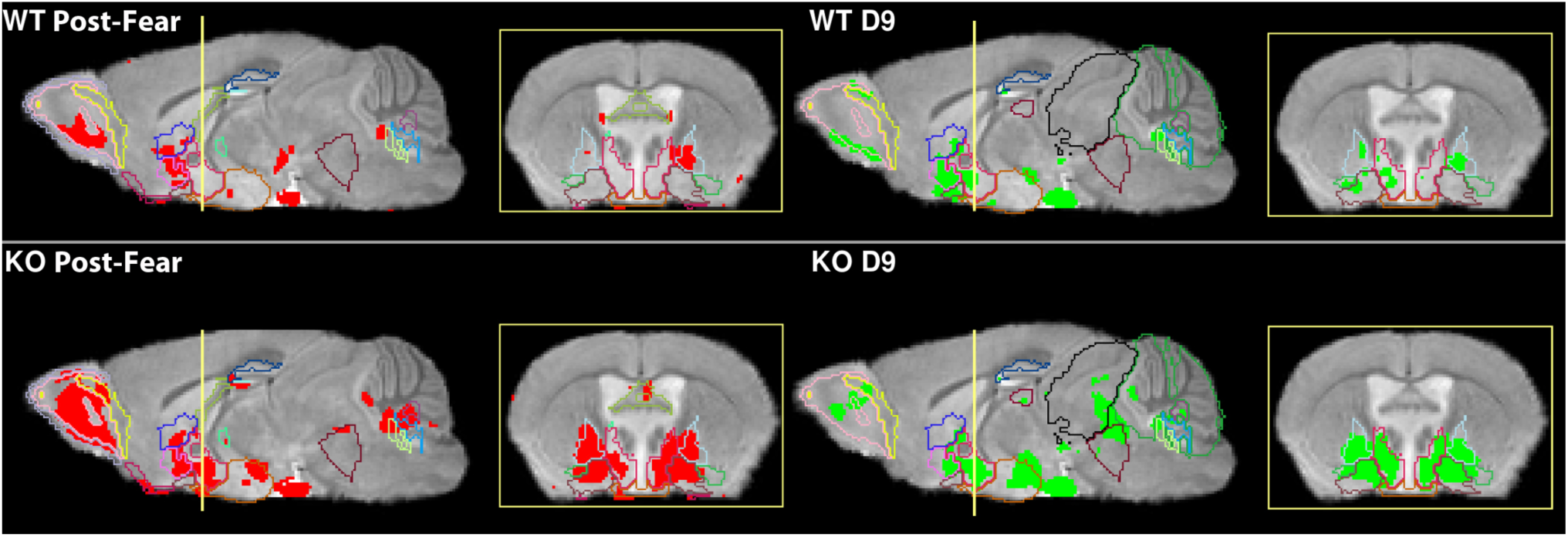
Enhanced regions extended beyond segment boundaries. **Left Panel**: Post-Fear enhanced regions (red) in WT (top) and in SERT-KO (bottom) in sagittal (bregma sagittal: ML -0.47mm) and corresponding coronal slices (bregma coronal: AP -0.5mm), at position indicated by a gold line on the sagittal slice. **Right Panel:** Day 9 activity (green) in WT (top) and SERT-KO (bottom). Slices are at the same location as in both panels, with active anatomical segments outlined. Note many regions where signal is contiguous across several segment boundaries, such as in SERT-KO ventral pallidum at both time points. All SPM maps at *p*<0.0001, *uncorr.* (T=5.4).

Although our imaging is voxel-wise and brain-wide, not all regions of the brain have yet been annotated in our *InVivo* Atlas, or in any other available MR atlases of living mouse brains to our knowledge. Our segmentation is based on an MRI atlas manually drawn on a Mn^2+^-enhanced MR image of a living mouse, in which insufficient intensity differences between sub-regions limited accurate identification. While we could guess at boundaries of these segments, we chose a more conservative approach for this version of our atlas and only annotated those regions whose outline could be definitively identified in Mn^2+^-enhanced MR images of a living mouse brain. Thus, in addition to signal in the annotated sub-regions described above, our SPM maps also contained significantly enhanced voxels in regions that were not annotated, and thus not included in our column graphs.

By comparing our activity maps with histological atlases, we can identify some of these non-annotated regions in specific slices (**Fig. 12**). For example, a strongly enhanced region between basolateral amygdala (BLA) and caudate putamen (CP) appears to correlate with central amygdala, not annotated in our atlas but significantly enhanced by SPM (**Fig. 12A**). In the brainstem the atlas is minimally segmented and thus our analysis may attribute activity to larger regions with many sub-regions not defined. Conversely, activity in small sub-regions within larger segments may not be included in our column graphs, as the large number of non-active voxels would drown signal in our ratiometric calculations. For example, in the pons, the signal in SERT-KO at Day 9 encircles the periaqueductal gray and overlays regions identified in the Allen Brain Atlas as the midbrain reticular nucleus (**Fig. 12B).** These regions were not originally sub-divided in the *InVivo* Atlas and are annotated as “midbrain” (MB), because their boundaries were not recognized in the MR image. Further work will be needed to identify and annotate these formerly invisible segments based on activity maps in live mice, thus building upon this high resolution *InVivo* Atlas with functional information for future applications.

**Fig. 12:**
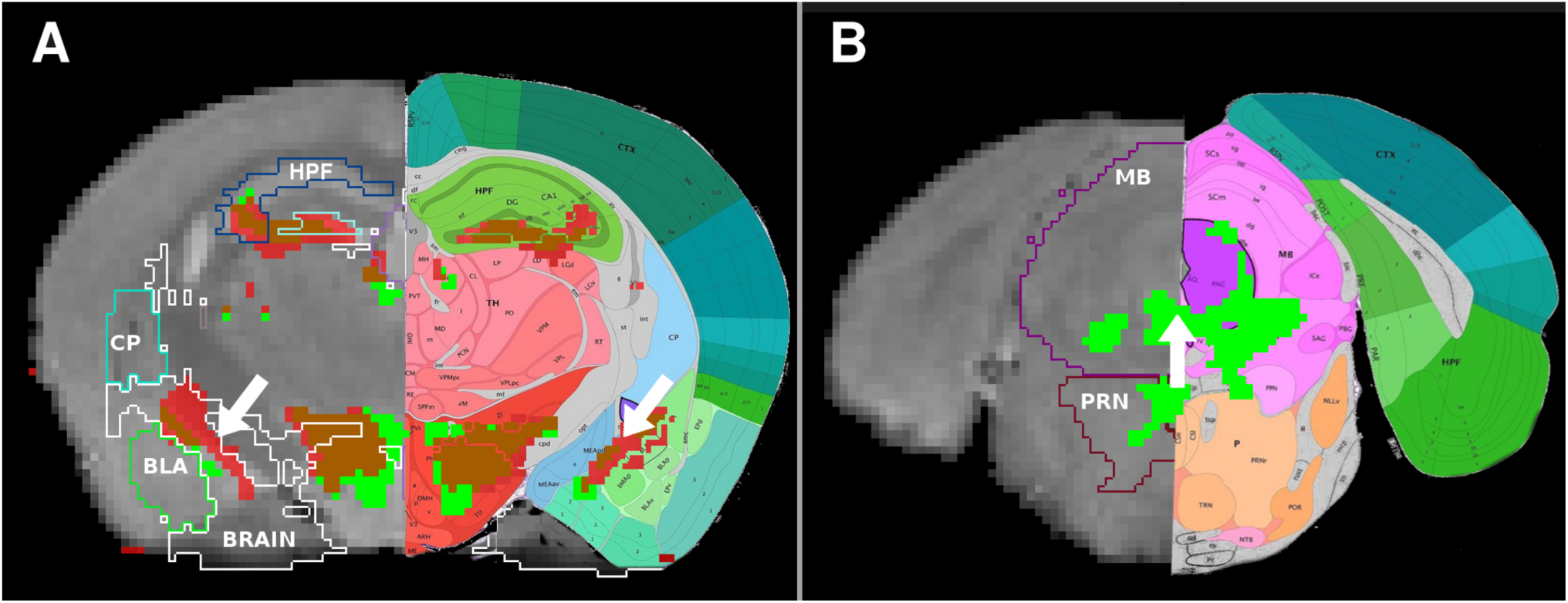
Activity was present in areas between annotated segments. Examples of MR slices with SPM activity in SERT-KO overlaid on our segmented gray scale template (left side) and on Mouse Brain Reference Atlas (right side of each slice), Allen Institute ^93^. Post-Fear (red), Day 9 (green), both (brown). **A**) Coronal slice through striatum (bregma coronal: AP -1.97mm) with white arrow indicating activity in region corresponding to the central amygdala in the Mouse Reference Atlas. “BRAIN” indicates non-segmented zones in the atlas. Abbreviations: hippocampal field (HPF); caudate-putamen (CP); basolateral amygdala (BLA); midbrain (MB); pontine reticular nucleus (PRN). **B**) Coronal slice through the midbrain (bregma coronal: AP -4.85mm) indicating activity in the periaqueductal gray (PAG), also not segmented in the *InVivo* Atlas. Regions are labeled as in the Allen Mouse Brain Reference Atlas ^93^.

## Discussion

Longitudinal whole brain MEMRI reveals evolution of regional neural activity from natural behavior, to fearful experience, to an anxiety state corresponding to exploratory or defensive behavior in living mice. We term these transitions “evolution” as changes develop gradually, with differing degrees and direction of change across regions from one state to the other over time. Neural activity transitions were not localized but occurred throughout the brain in 45 of the 87 sub-regions in our segmentation atlas. Some activity did not fluctuate but remained constant. For each state, the balance of activity between many segments differed, with some segments higher than others at one point and lower at another. Only with brain-wide measurements such as our MEMRI can this complex dynamic be witnessed.

In Pre-Fear images, after Mn^2+^ accumulation had occurred in freely moving mice, WT had minimal activity by SPM, while SERT-KO’s activity was high, with a pattern resembling WT Post-Fear. In Post-Fear images, in both genotypes, the olfactory system lit up, as expected from exposure to a strong novel odor. Also in Post-Fear, activity in both genotypes increased in many other brain regions, particularly in the striatum and ventral pallidum, hypothalamus and interpeduncular nucleus, which overshadowed small increases in the amygdala. SERT-KO sustained post-threat defensive behavior for 9 days while WT returned to normal exploration, both responses as expected ^3^. In tandem with behavior, SERT-KO brain activity remained high at Day 9 while WT transitioned to a new state, partly similar to its Pre-Fear. A striking difference in neural activity between genotypes at Day 9 was persistence of activity in SERT-KO striatum/pallidum, a region that projects to midbrain regulatory centers, as well as increased activity in the ventral tegmental area and sustained activity in thalamic nuclei, midbrain, and pons. Three regions stand out as having a much larger ratio of activity in SERT-KO than WT at Day 9: fundus of striatum (FS), dorsal raphe (DR) and pontine reticular nucleus (PRN), consistent with their roles in serotonin regulation and anxiety.

In this study, we analyze the consequences of a single fear event. Analysis of conditioned fear, where mice undergo multiple fearful events, has traditionally focused upon the amygdala ^110,111^. In fact the amygdala is not required for at least some types of innate fear responses in human ^112-114^. Animal studies of fear-related long-term potentiation in amygdala ^115^, lesion studies ^116^ and in humans with bilateral calcification within the amygdala who are unable to recognize fearful faces ^117,118^ further confirm a complex role of amygdala in fearful responses. Functional divisions within the central, medial, and basolateral amygdala determine the amygdala’s connections to structures in ventral pallidum in conditioned fear ^119-126^. The BST may be a first stop in a fear response, activating prior to the amygdala ^121,124,126^. However, conditioned fear is likely different from the single fear event studied here.

With brain-wide imaging, we find some activation in lateral amygdala in the immediate response to acute innate fear and activity in the central amygdala. This activation was not as great as in other brain regions, demonstrating that amygdala activation may not be the driver of an acute response to a single fear-provoking event ^127^. Our findings of activity in striatum and ventral pallidum, part of a network dubbed the “extended amygdala” ^128^, which increase at Day 9 are areas associated with anxiety ^129-131^. That we do not detect strong activity in basolateral amygdala immediately after PS may be because the amygdala activates later in the sequence of events following sensory perception of threat. Since we only image again 9 days later, our work does not determine whether or not the amygdala has activated and de-activated during that 9-day interval.

Evidence suggests a network of structures extending to and from the amygdala is involved in the evolution of fear to anxiety ^119,120,123,128,132^. The data we present here add to this work by witnessing activity in a set of regions in the anxious mouse, for example the fundus of striatum, bed nucleus of the stria terminalis, and the interpeduncular nucleus among others. With future mining of our data for network dynamics by repurposing emerging software from human MRI analyses, these network dynamics, and their connections/interactions, will be more precisely determined ^40,133^.

Our results of anxiety-like behavior corresponding to striatum/pallidum activity are supported by contemporary research in mouse and human. Until now most studies focus on a single region, hence speculations of how regions interact are based on aggregating multiple studies into a single composite ^103,134^. Inevitably, pre-selection of candidate regions provides a narrow view that emphasizes brain activity in one region while blind to all others. In addition when studies use different paradigms to elicit fear, results may appear inconsistent ^135-137^. Very recent reports of brain-wide electrophysiological recordings show a more comprehensive picture of sub-region interactions ^138^, although, like our atlas, these studies do not include all possible brain nuclei. Conceptualizing and mining rich datasets from whole brain imaging such as we report here obtained by MEMRI or by brain-wide electrode arrays is computationally challenging ^139^. By examining commonalities and discrepancies between studies, we may begin to ascertain neural representations of anxiety between mouse and human. In surveying the vast literature reporting studies on fear and anxiety, the specific type of fear provocation, and whether naturalistic or conditioned, likely plays a role in brain regions involved 134.

As neural activity evolves across the whole brain from acute fear-like to anxiety-like states, the balance of activation between segments changes. In Post-Fear images, the volume of active voxels increases in 19 different segments throughout the WT brain. Enduring “activity profiles” illustrate the persistence in some segments, while activity arising in new segments, or dissipating in others, reveals a dynamic progression, an evolution, from acute to prolonged defensive states. Our whole brain analysis shows that these states are not simply activation of a discrete segment but involve a dynamic balance of activity between segments. Progression of neural activity to Day 9, the anxiety state in SERT-KO, predominately evolves in the striatum and ventral pallidum, subcortical areas: targets of serotonergic neuromodulation and conserved across species 140,141.

Here we used the well-characterized SERT-KO mouse with disruption of the *SLC6α4* gene encoding SERT to generate an anxious phenotype ^13,142^, an extreme model of anxiety. This SERT-KO mouse has marked decrease in intracellular serotonin levels ^143-145^ and increased serotonin in interstitial fluid in the brain, with down-regulation of 5-HT_1A_ and 5-HT_1B_ receptor expression ^146-148^. These changes, likely a result of life long loss of SERT, are also found in humans after long-term treatment with selective serotonin reuptake inhibitors ^149^. SERT-KO mice have perturbed functional anatomy of prefrontal pyramidal cell projections ^69^, altered hypothalamic-pituitary-adrenal (HPA) axis output ^150^, and abnormal gut microbiome metabolism ^151,152^. The anxiety-like behavior of SERT-KO thus could be caused by a multitude of factors stemming from a deregulated serotonergic system both in the brain and throughout the body. We focused on quantifying functional brain-wide activity of the anxiety-like phenotype as an aggregate of all these influences, and not a specific response to PS. Anxiety-like behavior stemming from de-regulation of the serotonergic system may differ from anxiety-like behavior arising from other mechanisms.

Previous studies by optical or electrophysiological methods are hampered by the size of the brain, as localized field recording and imaging lack the ability to witness brain-wide activity or to penetrate deep structures. *In vivo* imaging by MEMRI provides an integrated longitudinal view of neural activity occurring throughout the brain in the awake behaving mouse ^16,17,19,26,31,47,56,57,64-66,68,153-158^. Corroborating evidence that MEMRI accurately represents neural activity includes activation of the olfactory system after odor with correspondence of MEMRI signal with expression of c-fos, an immediate early gene, which like Mn^2+^ also responds to neural activity via calcium currents ^44,159-161^. MEMRI is thus regarded as a reliable and powerful tool for imaging neural activity without the drawbacks of viral or synthetic agents that could alter endogenous neural activity in unpredictable ways.

We account for basal neural activity by imaging Mn^2+^ uptake occurring during 22-24h of freely behaving mice and comparing subsequent changes at successive times after predator odor in the same animals. This longitudinal strategy provides internal controls for the effects of manipulations, as each mouse serves as its own baseline. Since the WT mice return to baseline behavior at 9 days even with the most significant provocation (Mn^2+^-IP injection, exposure to predator odor, MR imaging), we did not attempt to perform parallel imaging at each time point in a separate set of mice to determine specific effects of each experience. It is possible that SERT-KO mice exposed only to handling and MR imaging would display prolonged anxious behavior without additional fear provocation. Hence we cannot specifically assign prolonged defensive behavior in SERT-KO *solely* to the predator odor experience. Our goal was to leverage the combination of a single predator stress provocation and SERT-KO genotype to produce mice with prolonged defensive behavior to compare their neural activity with a WT mouse experiencing similar conditions, and thereby identify neural correlates of that persistent defensive behavior. Without the predator stress, WT mice would not display defensive behavior, limiting their value as a normal control.

Although MEMRI allows for imaging *in vivo*, a drawback of MEMRI relative to optical imaging is less resolution due to low signal to noise ratio. MEMRI resolution can be increased with longer imaging times readily performed in fixed brains ^46^, although fixation precludes longitudinal studies. In living mice imaging longer than 2 hours requires extensive life support inside the scanner bore, technically challenging. In our study we obtained 100µm^3^ voxel resolution, encompassing ∼125 cells, in < 35 min scan time enabled by our high field MR scanner (11.7T). Another way to improve resolution and maintain statistical power would be to perform longer imaging sessions, although this is unlikely to reach better than a 50µm resolution and could further stress the animals.

For the first time, empowered by MEMRI and computational analysis that allows for automated co-registration and segmentation, we witness the evolution of neural activity between brain sub-regions, during and after a fearful experience. Imaging brain-wide activity in multiple animals over time produces a rich, complex dataset. We deploy our computational expertise to mine this dataset and find striking differences between time points and genotypes. However, quantifying the ratio of active to total voxels within each segment throughout the brain may under-estimate activity changes in larger regions with small areas of activity, and possibly over-estimate activity in smaller segments. Also, by aligning images of 12 individuals in each genotypic cohort, we miss individual differences, and by requiring voxels be within a 128-voxel cluster we also may miss small but significant tiny regions. Finally, our atlas does not segment all possible brain regions, and those not segmented are absent from this analysis. Activation by MEMRI or c-fos expression does not inform on whether regions are excitatory or inhibitory. In future experiments, brains fixed after MEMRI and imaged by whole brain light-sheet microscopy will provide the possibility of molecular identification of neuronal subtypes, whether stimulatory or inhibitory, to elucidate the function of activated segments, such as the medial pre-optic nucleus (MEP).

Our data reports brain regions activated by acute fear and the progression of brain-wide activity to the anxiety-state. A complimentary line of evidence on the role of the extended amygdala as a center of chronic anxiety adds to the interpretation of our results ^162^. We contribute to this literature by broadening the focus and revealing a brain-wide picture of dynamic activity throughout the brain, with many nuclei displaying greater activity than the amygdala, such as striatum and ventral pallidum, structures known to connect the amygdala to the hypothalamus. Through our statistical mapping, we uncover regions in the SERT-KO brain whose volume of activity is greater than WT nine days after an acute fear event, including paraventricular nucleus, ventral tegmental area, midbrain and pontine nuclei. Measuring the magnitude of intensity increases, we identify regions at Day 9 with 7-11% increased intensity in SERT-KO compared to WT, even in some regions statistically enhanced in both. Thus we uncover a dynamic balance of activities across and between many brain regions that together respond to acute fear and resolve or progress to anxiety states in a choreographed complex pattern. Changes in the balance of activity between structures rather than activity within any particular region appears to be involved in progression to the anxiety phenotype. Targeting these structures pharmacologically may disrupt the evolution from fearful responses to the anxiety brain state.

## Conclusions

MEMRI is a powerful tool that allows repeated *in vivo* imaging over time producing data to witness and quantify brain-wide activity as it evolves before, during and after experiences. Transition from natural behavior in the home environment to a single acute fear event culminating in persistent, chronic anxiety appears to depend on recruitment of not just one specific neural correlate but rather dynamic brain-wide networks between which the balance of activity evolves. In this paper we demonstrate that WT has little neural activity during normal exploratory behavior, responds to predator odor with increased activity in multiple brain segments and returns to baseline in behavior but not entirely by MEMRI by 9 days. In contrast, the SERT-KO mouse displays persistent defensive behavior in tandem with sustained or increased neural activity in multiple subcortical and brainstem structures where wild type activity declines. The areas with greatest differences between genotypes, such striatum and ventral pallidum, paraventricular nucleus, midbrain, dorsal raphe and pontine reticular nucleus, may represent neural correlates of anxiety. Thus, brain-wide activity mapping reported here reveals a complex dynamic with many brain regions participating in response to an acute fear experience and its resolution or progression to anxiety state. These results have the potential for translation to human fMRI results to inform future clinical research in drug targeting of sub-cortical and deeper brain structures.

## Acknowledgements

We thank Christopher S. Medina for guiding us in his processing protocols; Aaron Gonzales for pilot behavior sessions; Kathleen Kilpatrick for technical support with histologic preparation; Xiaowei Zhang for MRI session recordings; Sharon Wu Lin for animal staff support; and Kevin P. Reagan, Kyla Sorenson, Angela Miller and Daniel Perez for administrative assistance. We are grateful to Jonathan Brigman for helping set-up and perform the behavioral studies, Afonso Silva for critique of the protocol and the MS, and to Ralph Adolphs for helpful discussions on data analysis.

This research was supported in part by NIH NIMH RO1 MH096093 (EB), P50 GM085273 (EB) and the Harvey Family Endowment (EB). Additional support was provided by postdoctoral research fund ASERT NIH NIGMS 5K12GM088021(DB)

