## Supporting Information for "Evolution of brain-wide activity in the awake behaving mouse after acute fear by longitudinal manganese-enhanced MRI"

### Supporting information: Tables and Figures
### Uselman, Barto, Jacobs and Bearer

#### Supporting Information Figures & Legends

**Fig S1: Diagram of hypothesis on how fear may evolve to anxiety states.**

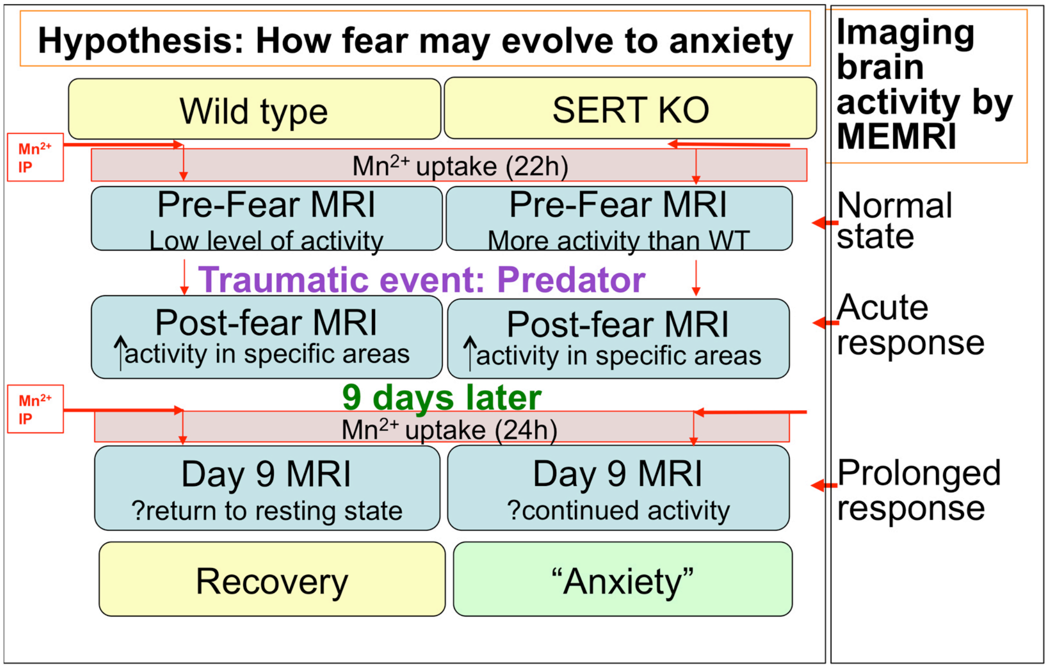


**Fig S1: Diagram of a hypothesis of how fear may evolve to anxiety states.** Wild type (WT) and SERT-KO mice differ in their ability to recover to baseline activity after a traumatic event (predator odor, PO). While WT recovers, SERT-KO remain in a defensive (anxiety) state. We hypothesize that this behavior reflects differences in neural activity in the brain, which we test directly with MEMRI brain-wide imaging. Horizontal red arrows indicate $Mn^{2+}$ IP injections, and vertical arrows in red boxes indicate $Mn^{2+}$ uptake into active neurons in naturally behaving, freely moving mice. Blue boxes represent three imaging sessions after $Mn^{2+}$ injection: Pre-Fear, Post-Fear and 9 days later (Day 9). Only one fear exposure is performed. We predicted that WT, with low neural activity prior to predator stress, responds to fear with increased activity, which resolves to Pre-Fear levels by 9 days afterwards. In contrast, SERT-KO has more neural activity prior to traumatic event, which increases in response to trauma. Unlike WT, activity in SERT-KO will not resolve at 9 days but continue and may evolve to new regions. This continued activity will manifest as anxiety-like defensive behavior in SERT-KO. In WT, some residual neural activity post-trauma may be present but at levels too low to produce an observable behavioral correlate.

**Fig.S2: Examples of individual images after skull-stripping (A) and averaged images of the datatsets after alignment (B)**

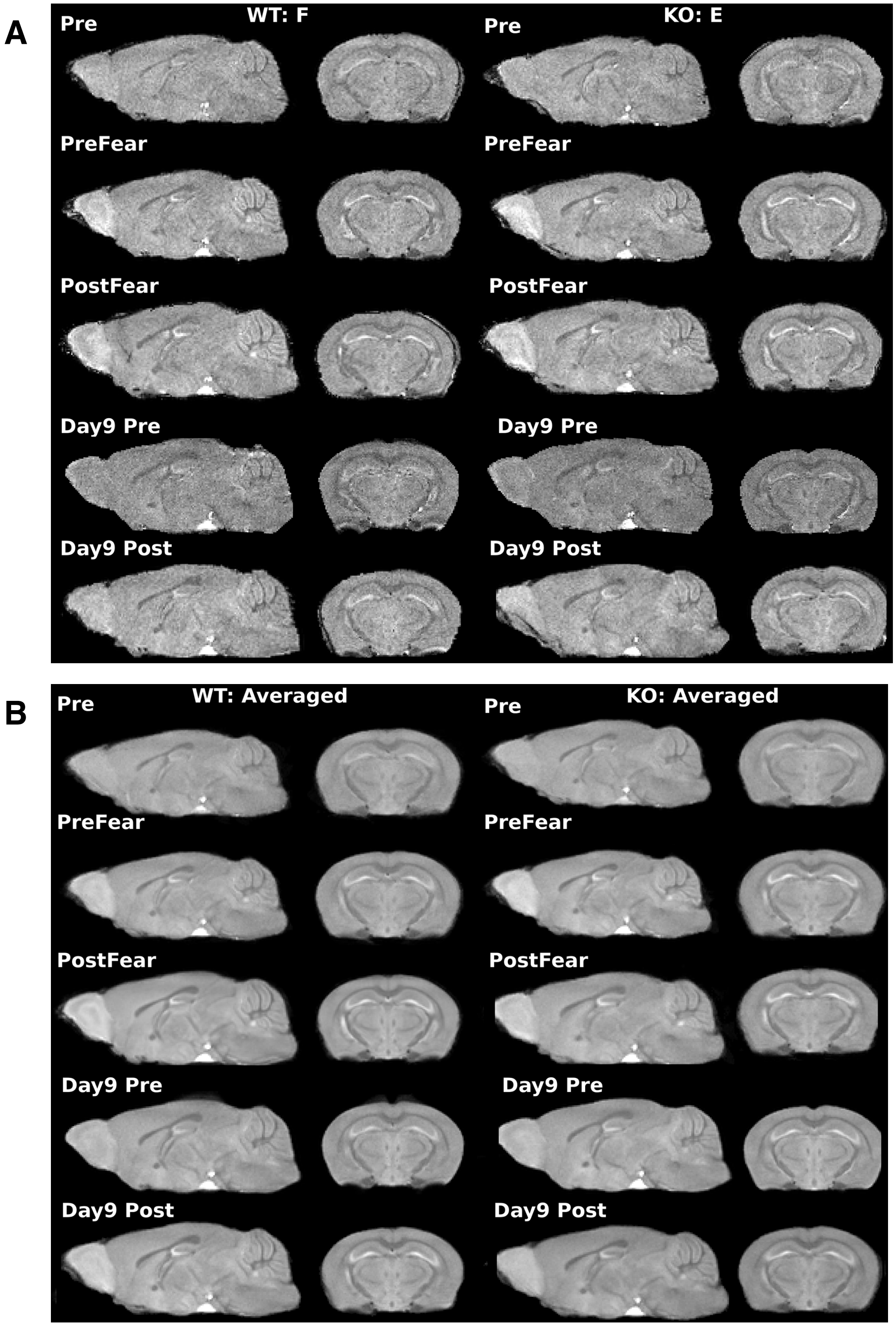


**Fig.S2: Examples of individual images after skull-stripping (A) and averaged images of the datatsets after alignment (B).** NOTE: Four image averaging of the **Post-Fear images (B) does not increase signal over single image (A)**

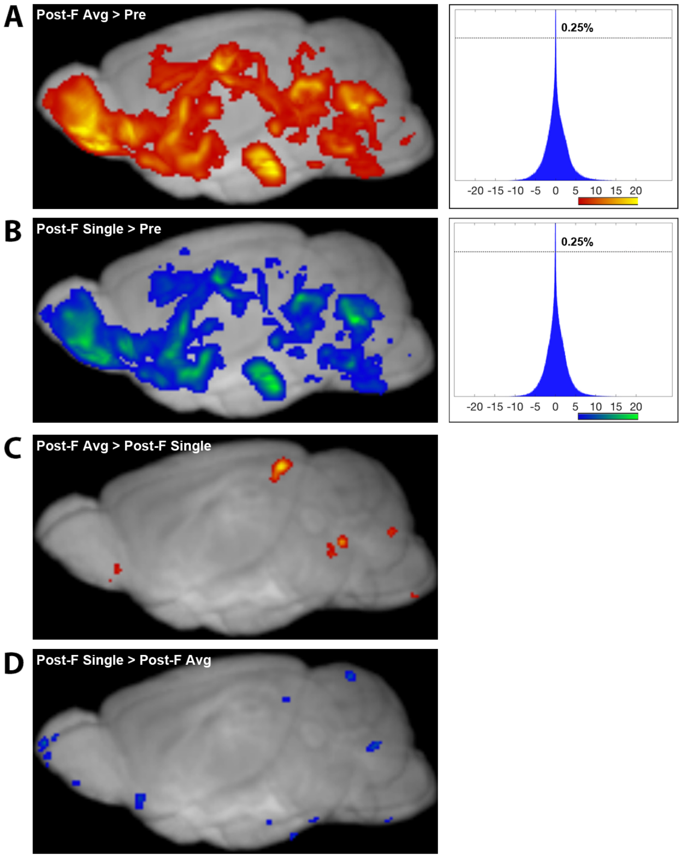


**Fig. S3: Averaging four Post-Fear images results in only very minor differences in SPM.**

**Rationale and Method:** To determine if averaging the series of four Post-Fear images improved enhanced statistically significant signal, we performed additional SPM comparisons between averages of 4 Post-Fear images and the fourth single Post-Fear image from that 4-image dataset (n=12). Two sets of pairwise t-tests were compared: (1) Averaged Post-Fear images (Post-F Avg>Pre) versus single Post-Fear images (Post-F 4>Pre), each to pre-injection images (**Fig. S3, A & B**); and (2) Single and averaged Post-Fear images compared in SPM to each other (**Fig. S3, C & D**). As before, we required a minimum voxel cluster size of 128 for significance at $p<0.0001$ uncorrected (T=5.4), the same statistic used for the other comparisons throughout this paper.

T-distribution histograms of SPM maps (**Fig. S3A** and **B**, right panels) confirmed the impression that averaging did not enhance sensitivity. If there were an increase in effect size and a decrease in variance produced by the averaging, we would see differences in the t-value histograms. Instead, 0.25% of the voxels in both single and averaged images reached the peak of the histogram, the variance was between -10 and +10 in both, and the ranges are approximately similar, -20 to +20.

Direct comparison of the single images to the averaged images detected only a few regions (**Fig. S3, C & D**). These regions likely represent those activated uniquely during the time frame of the fourth image and lost in the averaging procedure.

**<u>Figure Legend:</u>**

**Fig. S3: Averaging four Post-Fear images results in only minor differences in SPM.**

**A-B)** Snapshots of MRIcron 3D renderings and t-value histograms after paired t-tests in SPM contrasting Post-Fear images with pre-injection images. Results are overlaid on a gray scale template (left) and corresponding histogram of t-value distribution (right). **A)** Averaged Post-Fear images relative to pre-injection images (left) (red-yellow gradient according to t-value, see horizontal axis of the histogram for color-code). Corresponding t-value distribution is shown as a histogram (right). The line at the top of the graph labeled 0.25% indicates the percent of voxels at the mode. **B)** Single Post-Fear images relative to pre-injection images (left) (blue-green gradient according to t-value, see horizontal axis of the histogram for color-code). Corresponding t-value distribution is shown as a histogram (right) ($p<0.0001$, T=5.4).

**C-D)** Comparison between averaged Post-Fear images and single Post-Fear images. **C)** Averaged images with signal greater than single images (color code as in A). **D)** Averaged images with signal less than single images (color code as in B) ($p<0.0001$, T=5.4).

Conclusion: Only minor differences are found in SPM analyses between averaged Post-Fear images and a single image from the same dataset and the pre-injection image and between each other. Although these differences are small, they could be meaningful about progression of activity over the 2 hours immediately following a single fear exposure for future study. Hence, averaging is not responsible for the large amount of increased signal from Pre- to Post-Fear.

**Fig. S4. Horizontal column graphs across all time points for all segments in WT and SERT-KO.**

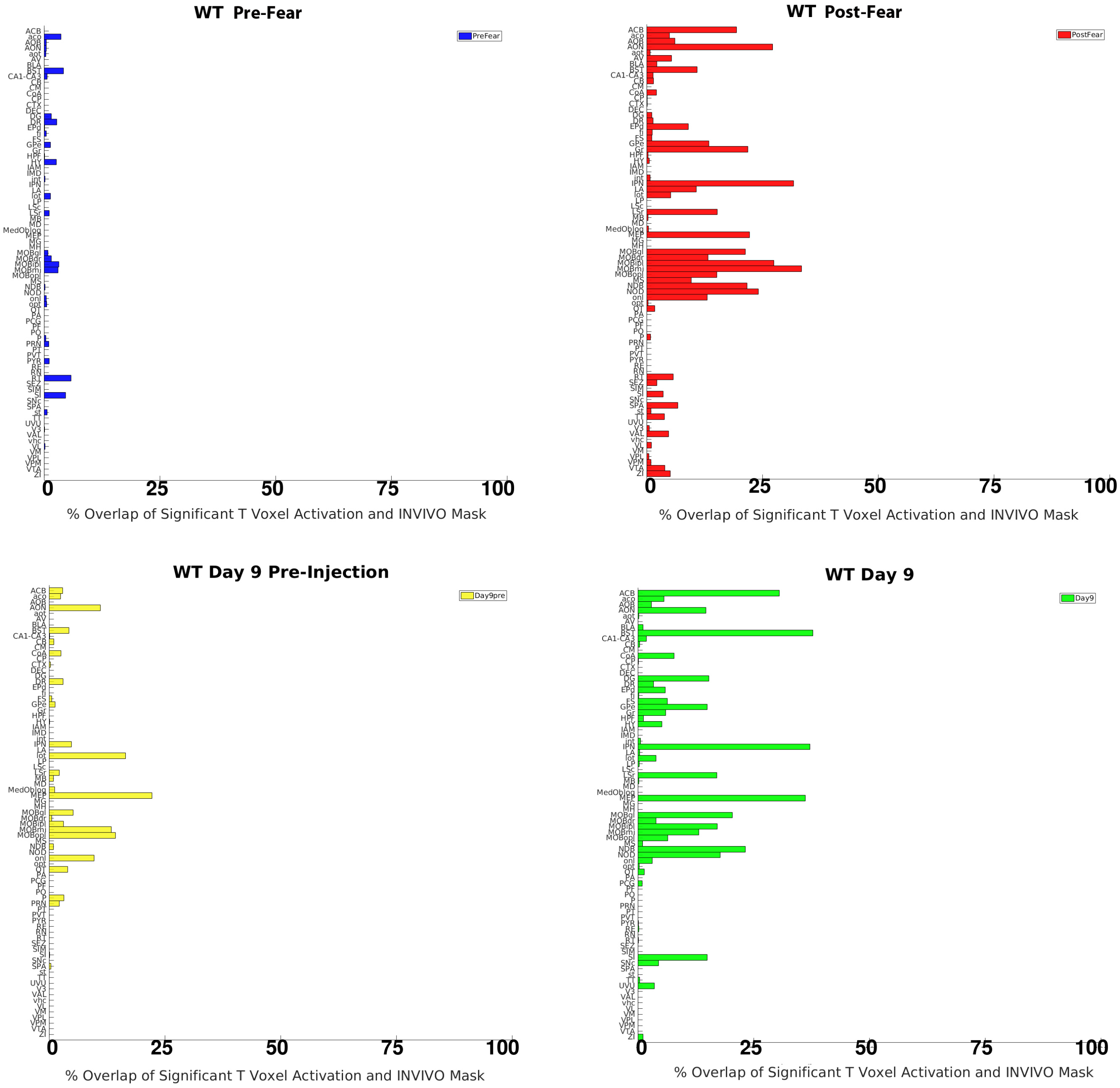


**Fig. S4A. WT.** Ratio of significant voxels/total voxels illustrates how activity transitions from Pre-Fear (blue) to Post-Fear (red), nine days later (yellow, Day 9 pre), finally to 9 days post fear (after the second $Mn^{2+}$ IP injection) (green, Day 9). Overall activity is sparse at Pre-Fear. Some small amount of signal is retained 9 days after the first $Mn^{2+}$ injection (Day 9 Pre). After the second $Mn^{2+}$ injection, (Day 9) some activity is sustained, some diminished and new areas turn on. Lack of signal in the Pre-Fear may be due to more random inconsistent neural activity across the 12 individuals, which would be lost in our statistical comparisons. A voxel-wise comparison of each individual animal with its respective pre-injection image may identify more individual-specific activity than is found with our statistical approach.

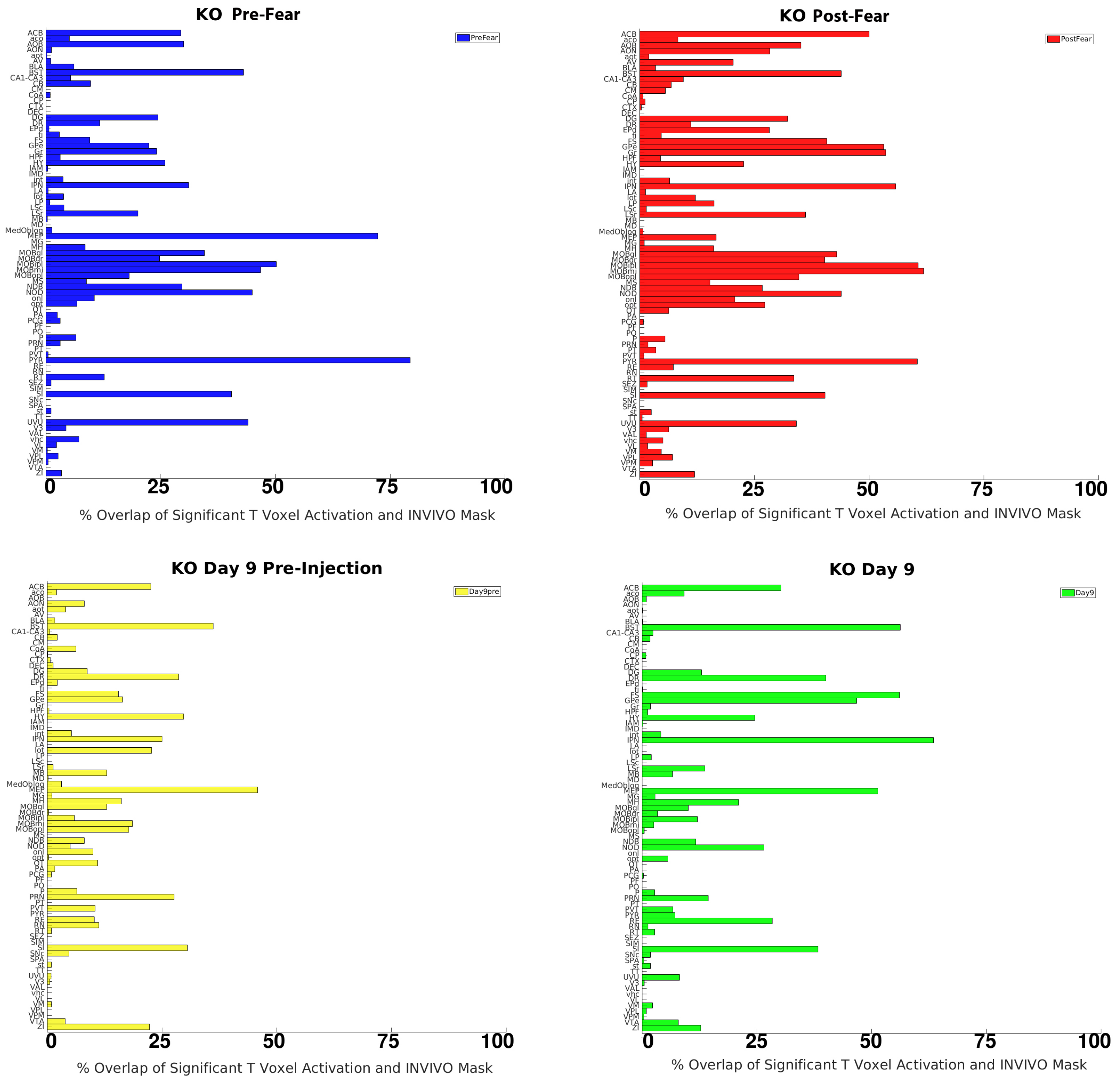


**Fig. S4B. SERT-KO.** Ratio of significant to total voxels illustrate how activity transitions from Pre-Fear (blue) to Post-Fear (red), to nine days later (yellow, Day 9 pre), finally to 9 days post $Mn^{2+}$ (green, Day 9) in SERTKO. Overall activity is much higher than WT and more signal is retained at Day 9 pre. New areas display signal at Day 9. The high amount of Pre-Fear activity in the vulnerable SERT-KO suggests that this mouse is already anxious at this time point, possibly reacting to the manipulations. Although differences between time spent in the light at Pre-Fear between WT and SERT-KO were not statistically significant (see **Fig. 4, main text**), a trend towards more defensive behavior was apparent. That we detect so much activity in the Pre-Fear image which reports an integrated composite of all neural activity occurring during the 22h post IP $Mn^{2+}$, when the mice are freely behaving in the home cage, suggests that the SERT-KO behavior is more stereotypical and uniform across individuals than WT, possibly a consequence of the disrupted neuromodulatory serotonergic system.

All 87 segments are shown. The order of segments is alphabetical, as in Table 1, which provides an alphabetized list of abbreviations.

**Fig. S5: A heat map showing how the ratio of active voxels compared to total voxels within each segment evolves over time.** In the heat map, segments are ranked according to the WT Pre-Fear analysis, from high (top) to low. (See **Supplemental Table S1** for abbreviations).

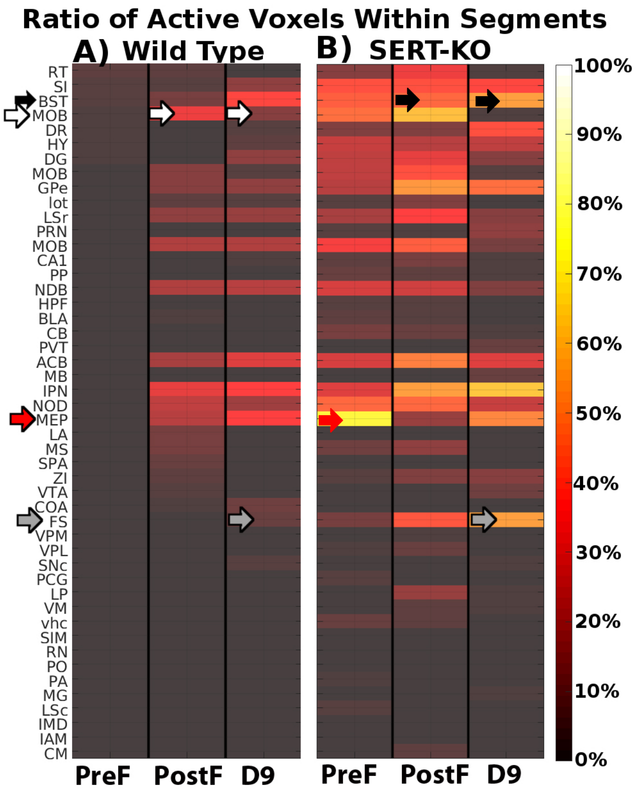


**A)** WT; **B)** SERT-KO. Total number of voxels in each segment were determined from the *InVivo* Atlas, and the number of active voxels within each segment obtained from segmented SPMs at each time point ($p$<0.0001 *uncorr*. T= 5.4) The heat map color code is indicated on the right. Calculations were preformed in a custom MATLAB script (see Material and Methods, main text).

The heat map shows little activity in the WT Pre-Fear (first column), with a larger number of segments in the SERT-KO both at Pre-Fear (fourth column) and post-Fear (fifth column). One segment stands out in the Pre-Fear data in SERT-KO: the median pre-optic nucleus (MEP) (see **Table 1** for abbreviations) (red arrow pointing to the yellow cell in the SERT-KO RS column). The MEP is a small nucleus in the hypothalamus with both inhibitory and excitatory connections throughout the brain in mouse and human. The heat map also demonstrates many other differences in specific regions across time points. For example, Post-Fear in both genotypes displays activity in numerous regions, with many more regions containing activity in SERT-KO than WT. Many of the brightest regions in the heat map in SERT-KO Post-Fear also have activity in WT but to a lesser extent. These areas included sensory systems, such as the main olfactory bulb, and association areas in the striatum, such as the diagonal band (NDB), interpeduncular nucleus (IPN) and globus pallidus (GP). One segment, the fundus of striatum (FS, gray arrow), displayed 41% active voxels in the SERT-KO. Of the 48 segments shown here, 31 displayed >5% increased volume of active voxels in the Post-Fear image for SERT-KO.

At Day 9 (D9) the WT has sustained, increased and decreased active voxels compared to Post-Fear. WT sustains some in the lateral septum, nucleus accumbens, and interpeduncular nucleus, and has 2-3 regions with increased activity such as the bed nuclei of the stria terminalis and substantia innominata. WT also displays decreases in the main olfactory bulb (MOB) and subparafascicular area (SPA), despite baseline exploratory non-defensive behavior in the light-dark box. SERT-KO at Day 9 has more active voxels than WT in the bed nuclei of the stria terminalis (BST) (black arrows) and substantia innominata (SI). SERT-KO accumulation in the main olfactory bulb (MOB, white arrows) is even further decreased than WT at D9 (third and sixth columns). SERT KO also show some increases at D9 over Post-Fear in the lateral amygdala (LA), bed nuclei of the stria terminalis (BST) and interpeduncular nucleus (IPN).

MEMRI detected increased activity compared to pre-Fear even though exploratory behavior had reverted to baseline exploratory behavior in WT, suggesting a neural signature for the memory of fear that is not strong enough to produce an observable behavioral effect. Indeed, it may be naive to believe that mouse brain activity always displays as a measureable behavior. Different ordering of these activity graphs reveal different information.

**Fig. S6. Between group comparison of WT and SERT-KO across all time points.**

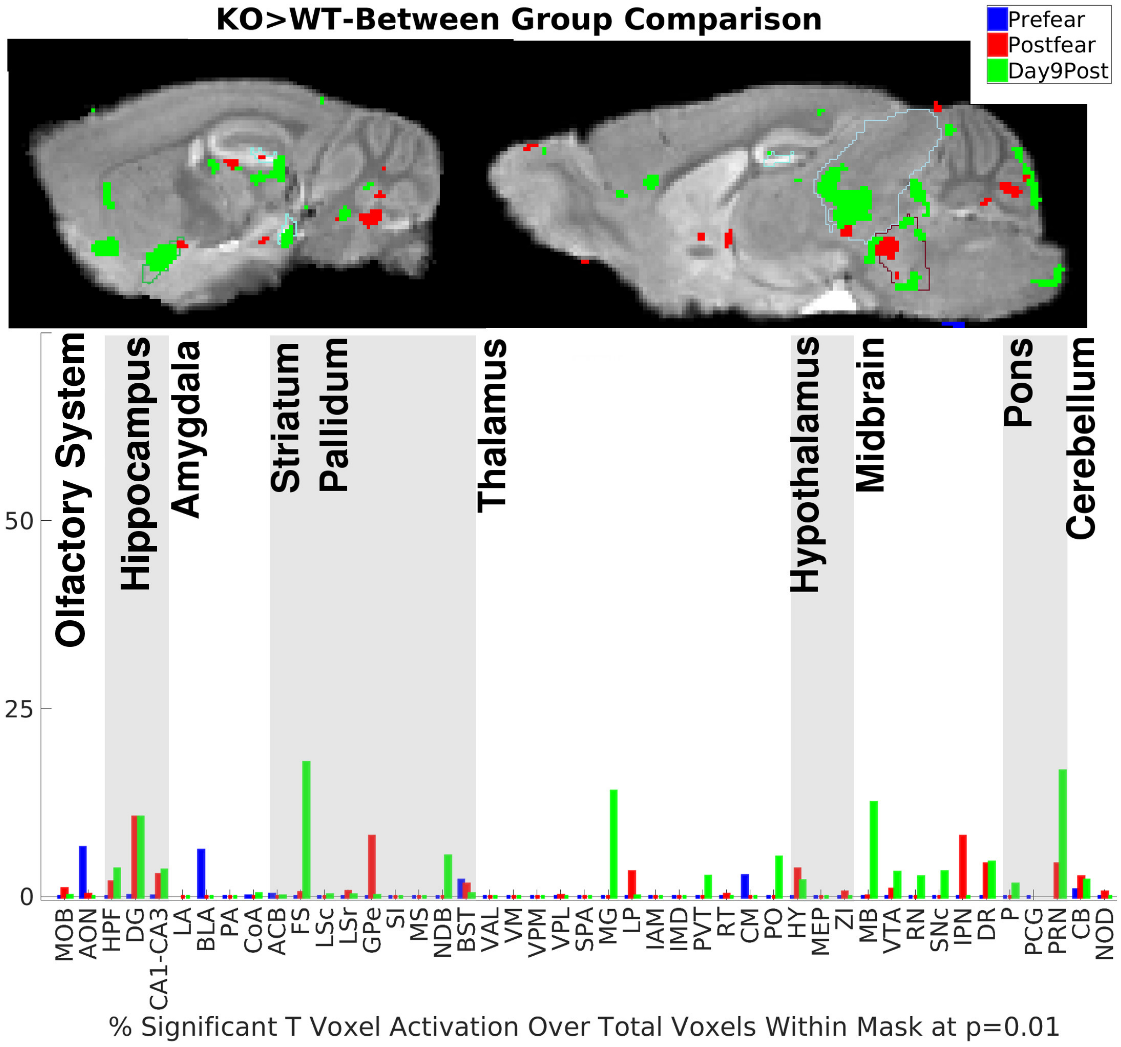


**Fig. S6. Between group comparisons of WT and SERT-KO across all time points.**

The SPM images from a two-sample t-test (top) identify regions where SERT-KO intensity increase is significantly greater than that of WT ($p$<0.01 *uncorr.*, T=2.7). Column graphs (bottom) show segments where the volume of significantly enhanced voxels is greater in SERT-KO than WT. Pre-Fear (blue); Post-Fear (red) and Day 9 pre (yellow) to Day 9 post $Mn^{2+}$IP (green). At Day 9, a greater volume of activity in SERT-KO appears in FS, NDB, MG, MB and PRN compared to WT. Our ROI analysis (**Fig. 9B**) demonstrated that the degree of difference in these areas at Day 9 was statistically significant as well. This analysis demonstrates that the major differences between genotypes are at Day 9 when behavior is also divergent.
