## Supplementary material for "Evolution of brain-wide activity in the awake behaving mouse after acute fear by longitudinal manganese-enhanced MRI": Video Caption

**S1. Video. Evolution of brain states for WT and SERT-KO**. This video depicts the progression of the SPMs from Pre-Fear (blue) to Post-Fear (red) and 9 days after the single predator stress fear exposure (Green). WT (left side) is displayed concurrently with SERT-KO (right side) to illustrate how the patterns of activity differ and evolve across the whole brain.
